# Mechanisms Driving Genome Reduction of a Novel *Roseobacter* Lineage Showing Vitamin B_12_ Auxotrophy

**DOI:** 10.1101/2021.01.15.426902

**Authors:** Xiaoyuan Feng, Xiao Chu, Yang Qian, Michael W. Henson, V. Celeste Lanclos, Fang Qin, Yanlin Zhao, J. Cameron Thrash, Haiwei Luo

## Abstract

Members of the marine *Roseobacter* group are key players in the global carbon and sulfur cycles. While over 300 species have been described, only 2% possess reduced genomes (mostly 3-3.5 Mbp) compared to an average roseobacter (>4 Mbp). These taxonomic minorities are phylogenetically diverse but form a Pelagic *Roseobacter* Cluster (PRC) at the genome content level. Here, we cultivated eight isolates constituting a novel *Roseobacter* lineage which we named ‘CHUG’. Metagenomic and metatranscriptomic read recruitment analyses showed that CHUG members were globally distributed and active in marine environments. CHUG members possess some of the smallest genomes (~2.52 Mb) among all known roseobacters, but they do not exhibit canonical features of genome streamlining like higher coding density or fewer paralogues and pseudogenes compared to their sister lineages. While CHUG members are clustered with traditional PRC members at the genome content level, they show important differences. Unlike other PRC members, neither the relative abundances of CHUG members nor their gene expression levels are correlated with chlorophyll a concentration across the global samples. Moreover, CHUG members cannot synthesize vitamin B_12_, a key metabolite made by most roseobacters but not by many phytoplankton species and thus thought to mediate the roseobacter-phytoplankton interactions. This combination of features is evidence for the hypothesis that CHUG members may have evolved a free-living lifestyle decoupled from phytoplankton. This ecological transition was accompanied by the loss of signature genes involved in roseobacter-phytoplankton symbiosis, suggesting that relaxation of purifying selection is likely an important driver of genome reduction in CHUG.

## Introduction

The marine *Roseobacter* group is a subfamily-level lineage in the *Alphaproteobacteria* and plays an important role in global carbon and sulfur cycling (1, 2). It is highly abundant in the coastal environments, accounting for up to 20% of all bacterial cells (3–5). Over 300 species and 100 genera have been described (6), the vast majority of which harbor large and variable genomes and grow readily on nutrient-rich solid media which are not representative of the niches found in the oligotrophic oceans. Early culture-independent 16S rRNA gene surveys showed that the oceanic roseobacters are represented by a few uncultivated lineages (1, 7). Recently, novel cultivation techniques and single-cell genomics have made available (partial) genome sequences of several previously uncultivated lineages including NAC11-7 (8), DC5-80-3 (also called RCA) (9, 10) and CHAB-I-5 (11, 12). Although these lineages are spottily distributed throughout the *Roseobacter* phylogeny, they together form a pelagic *Roseobacter* cluster (PRC). The PRC members consistently harbor smaller genomes and show more similar genome content compared to other roseobacters (11). Learning their evolutionary histories is essential to understand how the genetic and metabolic diversity of the pelagic *Roseobacter* lineages has formed, which in turn helps appreciate their roles in oceanic carbon and sulfur cycles. However, most PRC members form orphan branches and lack closely related reference genomes, which hampers our further understanding of their evolutionary trajectories.

Here, we isolated eight closely related roseobacters from several ocean regions that consistently possessed some of the smallest genomes (~2.52 Mb) among all known roseobacters. They together formed a novel *Roseobacter* lineage which we named ‘CHUG’ (Clade Hidden and Underappreciated Globally) that was abundant and active in global oceans. Unlike other PRC members, the global distribution of CHUG members was uncorrelated with chlorophyll a (Chl-a) concentration and they cannot *de novo* synthesize vitamin B_12_, which is often the metabolite roseobacters supply to phytoplankton during their symbiosis (2, 13–15). Therefore, the reductive evolution of CHUG may also indicate a dissociation with phytoplankton, a feature so far unique to CHUG among pelagic roseobacters.

## Materials and Methods

Detailed methods are described in Supplementary Text 1. Briefly, samples were collected from surface water of the South China Sea, the East China Sea and the northern Gulf of Mexico. Over 20 CHUG isolates were retrieved following different dilution cultivation procedures, and genomes of eight isolates from the three ocean regions were sequenced with Illumina platforms, assembled with SPAdes (16) and annotated with Prokka (17). Among these, the isolate HKCCA1288 was further sequenced with PacBio Sequel platform to obtain a complete and closed genome. The average nucleotide identity (ANI) between genomes was calculated using fastANI (18). The assembled genome size, gene number, coding density and GC content of each genome were obtained using CheckM (19), whereas the estimated genome size was adjusted as (*assembled genome size*)/(*completeness*[+[*contamination*) (20). Pseudogenes were predicted following our recent study (21), and other genomic features were summarized using custom scripts (https://github.com/luolab-cuhk/CHUG-genome-reduction-project). The phylogenetic ANOVA analyses were performed to compare the analyzed genomic traits while controlling for the evolutionary history of those traits using the ‘phylANOVA’ function of the ‘phytools’ R package (22).

The *TARA* Ocean metagenomic and metatranscriptomic sequencing data with size fractions up to 3 μm (prokaryote-enriched) (23, 24) and metagenomic sequencing data with size fraction of 5-20 μm (nanoplankton-enriched) (25) were mapped to all 79 roseobacters studied here using bowtie (26) and BLASTN (27). Only reads sharing >95% similarity and >80% alignment to their best hit were kept for relative abundance and activity calculation. The correlation analysis was performed using the ‘rcorr’ function in the ‘Hmisc’ R package (28), and the significance level was adjusted using stringent Bonferroni correction for multiple comparisons.

The *Roseobacter* phylogeny was constructed based on 120 bacterial marker genes (29), and the reference *Roseobacter* genomes included in the phylogeny followed a previous study (30). The orthologous gene families were predicted with OrthoFinder (31), and a binary matrix of the presence and absence pattern of orthologous gene families were used to construct the genome content dendrogram. The gene copy number of each orthologous family was further used to estimate the ancestral genome content for CHUG, its sister group and the outgroup using BadiRate (32).

## Results

### The CHUG diversity

Eight strains constituting a novel lineage (Fig. 1A) within the *Roseobacter* group, which we named Clade Hidden and Underappreciated Globally (CHUG), were isolated from the coastal waters of the South China Sea, the East China Sea, and the northern Gulf of Mexico (Table S1). Their genomes shared ≥99.7% 16S rRNA gene identity and ≥93% whole-genome average nucleotide identity (ANI). The CHUG lineage further exhibited ≥98.2% 16S rRNA gene identity when sequences of a few uncultivated members were added (Fig. S1), which was comparable to other pelagic *Roseobacter* lineages (98% (10) for DC5-80-3 and 96% (33) or 98% (7) for CHAB-I-5). CHUG genomes had ≤96.5% 16S rRNA gene identity and ≤71% ANI compared to the sister group members (Fig. 1A). All CHUG isolates were sampled exclusively from pelagic environments, whereas members of their sister group and the outgroup inhabit highly diverse salty environments including pelagic ocean, saline lake, algal culture and coastal sediment (Table S1).

**Fig. 1.**
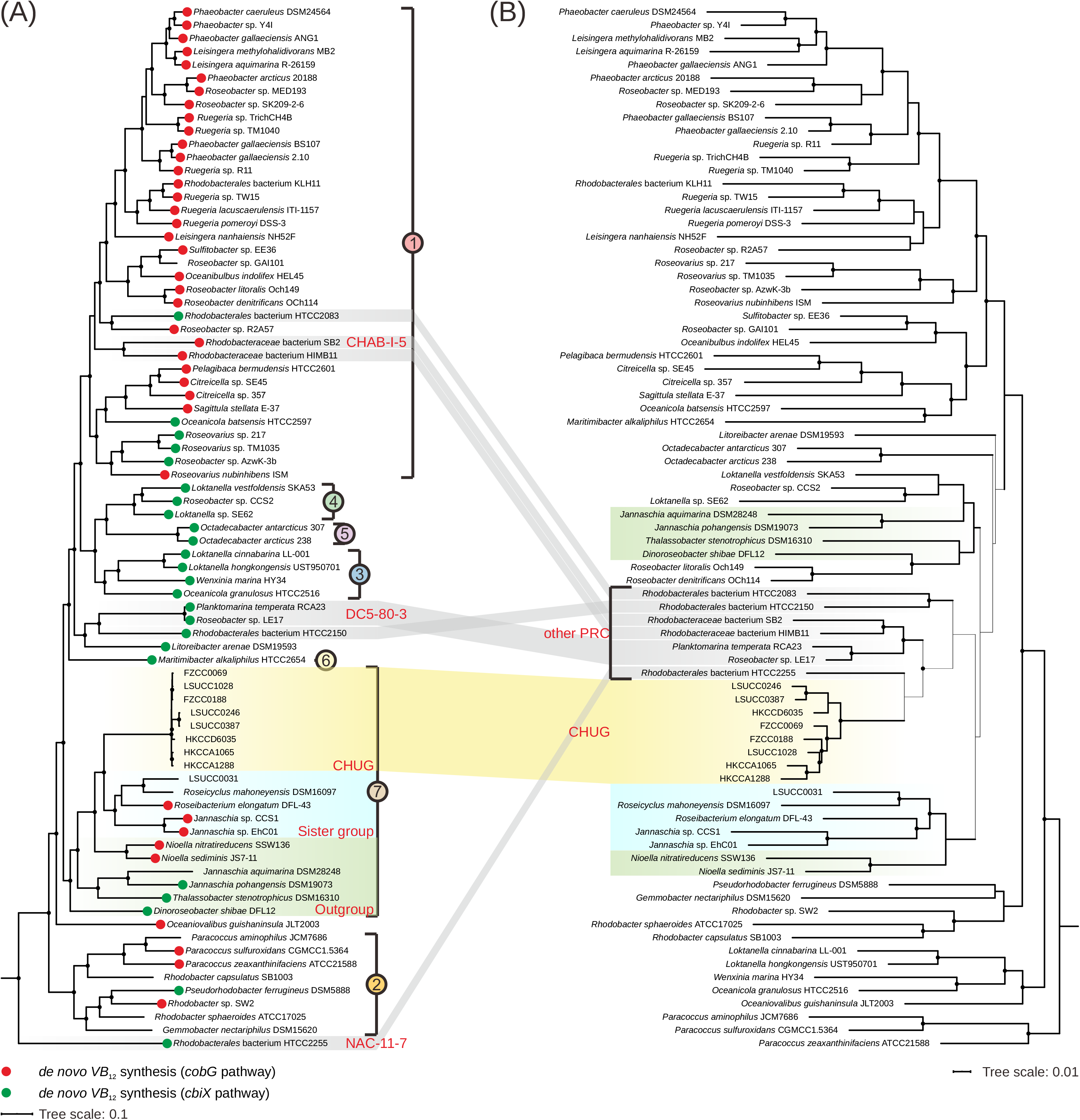
**(A)** Maximum likelihood phylogenomic tree showing the position of CHUG in the *Roseobacter* group. The phylogeny was inferred using IQ-TREE (91) based on a concatenation of 45,904 amino acid sites over 120 conserved bacterial proteins (29). Solid circles in the phylogeny indicate nodes with bootstrap values >95%. The potential of aerobic (key gene *cobG*, red) and anaerobic (key gene *cbiX*, green) cobinamide synthesis (the first stage of Vitamin B_12_ synthesis) is labeled at the tips. Subclades of the *Roseobacter* group are marked according to a recent study (30). **(B)** Dendrogram of the same *Roseobacter* genomes based on the presence/absence pattern of orthologous gene families.

We also constructed a dendrogram based on the presence/absence pattern of orthologous gene families (Fig. 1B). Although not monophyletic in the phylogeny based on shared genes (Fig. 1A), CHUG and seven other genomes from taxa previously sampled from pelagic environments formed a coherent group called the Pelagic *Roseobacter* Cluster (PRC) (11). One previously identified PRC member, *Roseobacter* sp. R2A57 (4.13 Mb), was not affiliated with PRC in this study. To facilitate our analysis and discussion, we divided the 79 roseobacters used in the present study into five groups: CHUG (eight genomes), its sister group (five genomes), the outgroup of CHUG and its sister group (six genomes), other PRC members (seven genomes) and other reference roseobacters (53 genomes).

### Genomic features

Among the eight CHUG genomes, one (HKCCA1288) was closed with 2.66 Mb and the remaining draft genomes were nearly complete (≥98.5%) according to CheckM predictions (Table S1). Among other roseobacter genomes under comparison, at least 17 genomes were closed and the remaining ones were nearly complete (≥96.5%) (Table S1). Based on the assembled genome sizes, CHUG members possessed much smaller genomes (2.52 ± 0.07 Mb, Fig. 2A) than an average roseobacter (4.16 ± 0.68 Mb). Further, their genome sizes were comparable to those of the NAC11-7 cluster represented by the strain HTCC2255 (estimated complete size to be 2.34 Mb), which is a basal roseobacter with the smallest genome among all known roseobacters (34). As in HTCC2255, no plasmids were found in the CHUG genomes. However, the coding density of CHUG (91.7 ± 0.5%, Fig. 2B) showed no significant difference with its sister group and the outgroup (90.7 ± 0.7% and 90.7 ± 0.6%, respectively) based on the phylogenetic ANOVA analysis (*p* > 0.05, ‘phylANOVA’ in the ‘phytools’ R package; the same test used below unless stated otherwise), which performs a simulation-based comparison while taking into account the influence of phylogeny on the trait evolution (22). CHUG genomes had a lower genomic GC content (55.4 ± 0.8%, Fig. 2C) compared to their sister group (63.5 ± 1.6%, *p* < 0.05), although no significant difference was identified compared to the outgroup (63.8 ± 2.6%). In terms of pseudogenes, the number (99 ± 24, Fig. 2D) and ratio (3.9 ± 0.9%, Fig. 2E) in CHUG members were not significantly different from those of the sister group (128 ± 51; 3.3 ± 1.1%) and outgroup (148 ± 37; 3.7 ± 0.9%). The seven other PRC members also had smaller genomes (3.26 ± 0.51 Mb, Fig. 2A) and a reduced GC content (49.6 ± 5.5%, Fig. 2C) compared to the 53 other reference roseobacters (genome size: 4.32 ± 0.64 Mb, GC content: 61.9 ± 4.1%; *p* < 0.01), but there was no significant difference between the two groups in terms of the coding density (90.4 ± 0.9% for seven PRC members versus 89.3 ± 1.5% for other roseobacters, Fig. 2B), or the number (108 ± 49 for seven PRC members versus 205 ± 134 for other roseobacters, Fig. 2D) and ratio of pseudogenes (3.2 ± 1.4% for seven PRC members versus 4.7 ± 2.4% for other roseobacters, Fig. 2E).

**Fig. 2.**
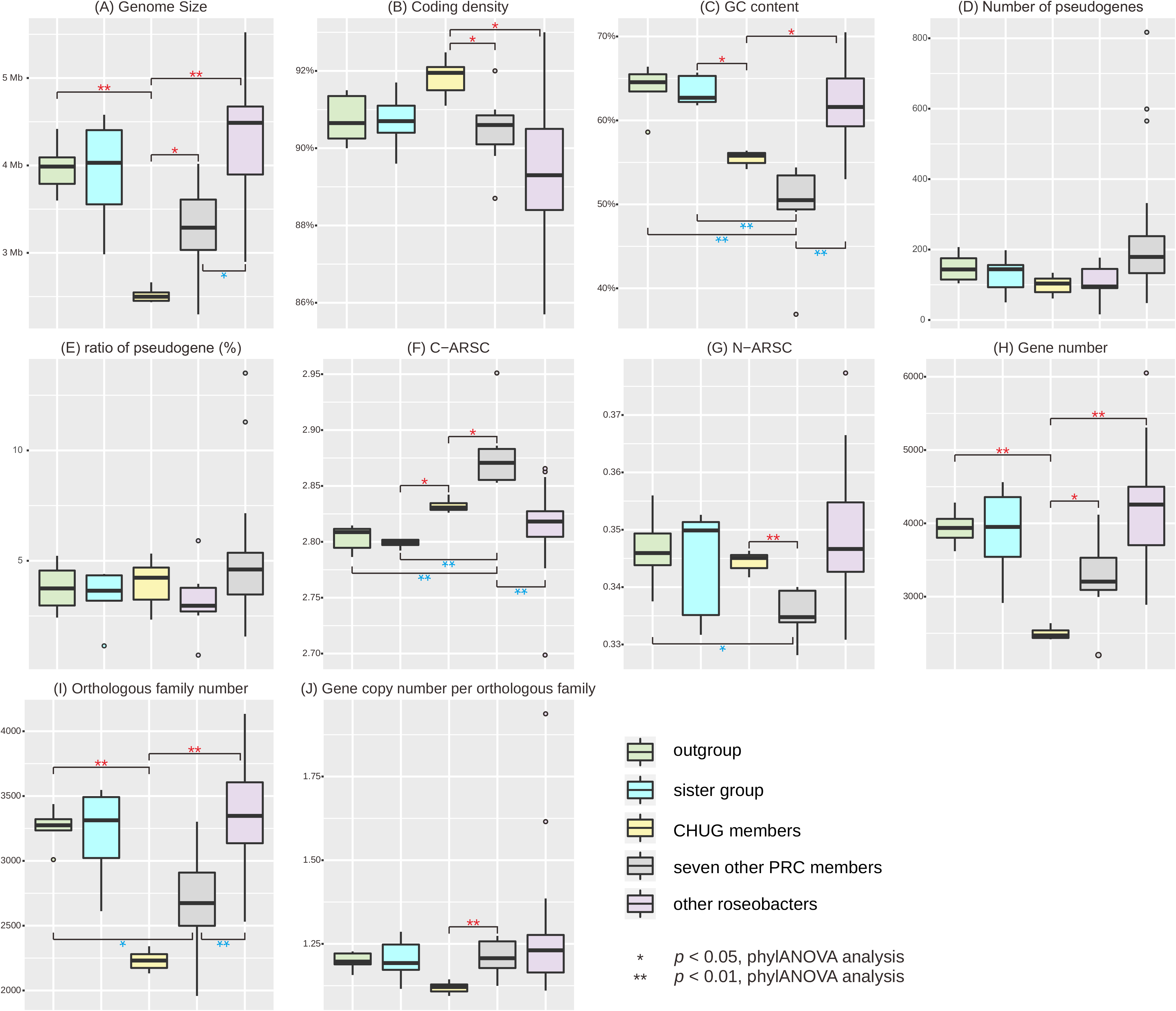
Genomic feature comparisons between CHUG, their sister group, the outgroup, seven other PRC members, and other reference roseobacters. The significance level in genomic features between CHUG and the other four groups are shown in red, while that between seven other PRC members and the remaining groups are shown in blue. The markers * and ** denote *p* < 0.05 and *p* < 0.01 (phylANOVA analysis (22)), respectively. Abbreviations: C-ARSC, carbon atoms per amino-acid-residue side chain; N-ARSC, nitrogen atoms per amino-acid-residue side chain.

CHUG genomes showed increased use of carbon atoms per amino-acid-residue side chain (C-ARSC, 2.833 ± 0.005, Fig. 2F) compared to the sister group (2.799 ± 0.004, *p* < 0.05). However, no significant difference was found in CHUG members in the use of C-ARSC compared to the outgroup (2.803 ± 0.011), nor that of nitrogen atoms per amino-acid-residue side chain (N-ARSC, 0.345 ± 0.002, Fig. 2G) compared to the sister group (0.344 ± 0.008) or the outgroup (0.346 ± 0.006). Likewise, the seven other PRC genomes had significantly higher C-ARSC (2.879 ± 0.031, Fig. 2F) than the 53 other reference roseobacters (2.817 ± 0.026, *p* < 0.01), but there was no significant difference between their N-ARSC (0.336 ± 0.004 for seven PRC members versus 0.348 ± 0.009 for other roseobacters, Fig. 2G).

We further investigated the codon usage and amino acid usage patterns in these lineages. The CHUG genomes tended to comprise codons with more adenine/thymine (A/T) and less guanine/cytosine (G/C) for 11 amino acids compared to the sister group and for 12 amino acids compared to the outgroup, respectively (*p* < 0.05, Fig. S2 and Supplementary Text 2.1). Furthermore, CHUG members possessed a higher fraction of isoleucine and lysine in their proteomes but a lower fraction of glycine, proline, valine and tryptophan when compared to the sister group or outgroup (*p* < 0.05, Fig. S3), which may be partially affected by the differences of nitrogen (N) use in their corresponding codons (Supplementary Text 2.1).

Consistent with their genome size differences, CHUG genomes contained a significantly smaller number of coding genes (2,486 ± 78, Fig. 2H) compared to the outgroup (3,939 ± 214, *p* < 0.01) and the seven other PRC genomes (3,253 ± 545 genes, *p* < 0.05). The CHUG genomes contained 2,215 ± 70 orthologous gene families (Fig. 2I) with 1.12 ± 0.01 gene copy per family (Fig. 2J). By comparison, the outgroup genomes contained 3,259 ± 130 families (*p* < 0.01) orthologous gene families with 1.20 ± 0.04 (*p* > 0.05) gene copy per family, while the seven other PRC genomes possessed 2,678 ± 398 families (*p* > 0.05) orthologous gene families with 1.21 ± 0.05 (*p* < 0.01) gene copy per family. No significant difference occurred between CHUG and the sister group (3,865 ± 591 genes, 3,197 ± 345 gene families and 1.20 ± 0.05 copy per family). Additionally, while the number of genes and number of gene copies per family of the seven other PRC genomes was not significantly different from those in the 53 other reference roseobacters (4,199 ± 644 genes and 1.25 ± 0.12 copy per family, Fig. 2H,J), the seven other PRC genomes had fewer orthologous families compared to the 53 other reference roseobacters (3,362 ± 362, *p* < 0.01, Fig. 2I).

### Global distribution and ecological drivers

We used recruitment analysis with the global-scale *TARA* Ocean metagenomic and metatranscriptomic datasets with size fractions up to 3 μm (prokaryote-enriched) (23, 24) to quantify the global distribution of CHUG and other PRC members. The eight CHUG members recruited 0.0005% and 0.0008% of all metagenomic (Fig. 3A) and metatranscriptomic (Fig. 3B) reads, respectively. The CHUG members appeared to be less abundant and less active than other PRC representatives such as *Rhodobacterales* bacterium HTCC2255 (NAC11-7), *Rhodobacteraceae* bacterium SB2 (CHAB-I-5) and *Planktomarina temperata* RCA23 (RCA or DC5-80-3) (Welch’s t-test, *p* < 0.01 for each). A similar pattern was also found using *TARA* Ocean metagenomic sequencing data with the size fraction of 5-20 μm (nanoplankton-enriched; Fig. 3C) (25). The CHUG members further represented 1.165% of the total reads from the nutrient perturbation experiments in mesocosm situated in the Red Sea (Fig. S4A) (35), and they also showed seasonality, as they recruited 0.007%, 0.032% and 1.623% of the total reads sampled at Kwangyang bay ocean (36) in February, May and August 2015, respectively (Fig. S4B).

**Fig. 3.**
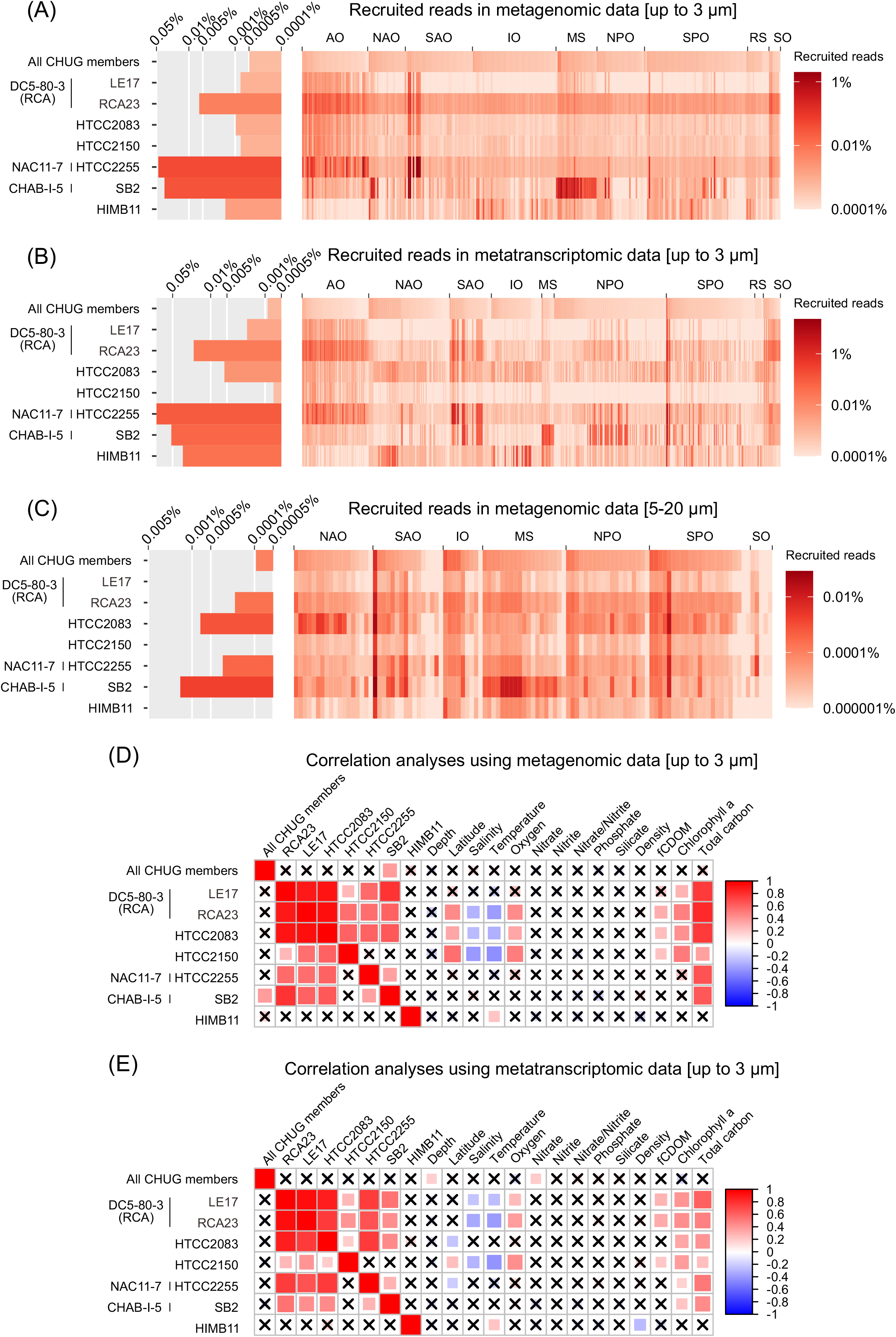
The global distribution of CHUG and its ecological correlation with environmental factors. **(A, B & C)** The relative abundance of CHUG and other PRC members in the bacterial communities based on recruitment analysis using the metagenomic *TARA* Ocean sequencing samples with size fractions up to 3 μm (A), and metatranscriptomic sequencing samples with size fractions up to 3 μm (B), and metagenomic sequencing samples with size fraction of 5-20 μm (C). **(D & E)** Correlation analysis between the relative abundance of CHUG and other PRC members and environmental parameters measured in the *TARA* Ocean metagenomic (D) and metatranscriptomic (E) samples. The *p* value is adjusted using stringent Bonferroni correction. Nonsignificant correlations are indicated by crosses for *p* > 0.05 after adjusting. Abbreviations: AO, Arctic Ocean; NAO, North Atlantic Ocean; SAO, South Atlantic Ocean; IO, Indian Ocean; MS, Mediterranean Sea; NPO, North Pacific Ocean; SPO, South Pacific Ocean; RS, Red Sea; SO, Southern Ocean; fCDOM, fluorescence, colored dissolved organic matter.

Next, we sought to identify the ecological factors that may drive the global distribution and activity of the CHUG members, and to compare it to seven other PRC members using the *TARA* Ocean metagenomic (Fig. 3D) and metatranscriptomic (Fig. 3E) samples. The relative abundance and activity of CHUG members and the PRC member *Rhodobacteraceae* bacterium HIMB11 were not correlated with other PRC members, chlorophyll a (Chl-a) concentration, or the total carbon (Fig. 3D,E; Bonferroni corrected *p* < 0.05). On the other hand, the relative abundances of other PRC members were mutually positively correlated with each other, with Chl-a concentration, and with total carbon in both metagenomic and metatranscriptomic samples (Fig. 3D,E; Bonferroni corrected *p* < 0.05). In addition, the activity of CHUG genomes was positively correlated with nitrate and depth (Fig. 3E; Bonferroni corrected *p* < 0.05). From a gene-centric perspective, 58.6% ± 1.2% and 88.3% ± 12.7% genes from the eight CHUG genomes and seven other PRC members recruited *TARA* metatranscriptomic reads, respectively. Among the most expressed gene families (top 5%), many were housekeeping genes involved in transcription, translation, cell cycle, respiration, the tricarboxylic acid cycle (TCA) cycle, and the biosynthesis of amino acids, chaperones, cell wall, and capsule (Fig. 4). Both CHUG and other PRC members also had highly expressed genes for light utilization (e.g. the photosynthesis gene cluster or proteorhodopsin) and nutrient (e.g. carbohydrates and amino acid) transport. Additional highly expressed genes among CHUG members included those involved in zinc transport, the cytochrome *cbb_3_*-type oxidase, acetate transporters, and genes for mercury homeostasis, among which the latter two were exclusively found in CHUG members (Fig. 4). On the other hand, some highly expressed orthologous gene families specific to the seven other PRC members were related to phosphonate transport and taurine degradation.

**Fig. 4.**
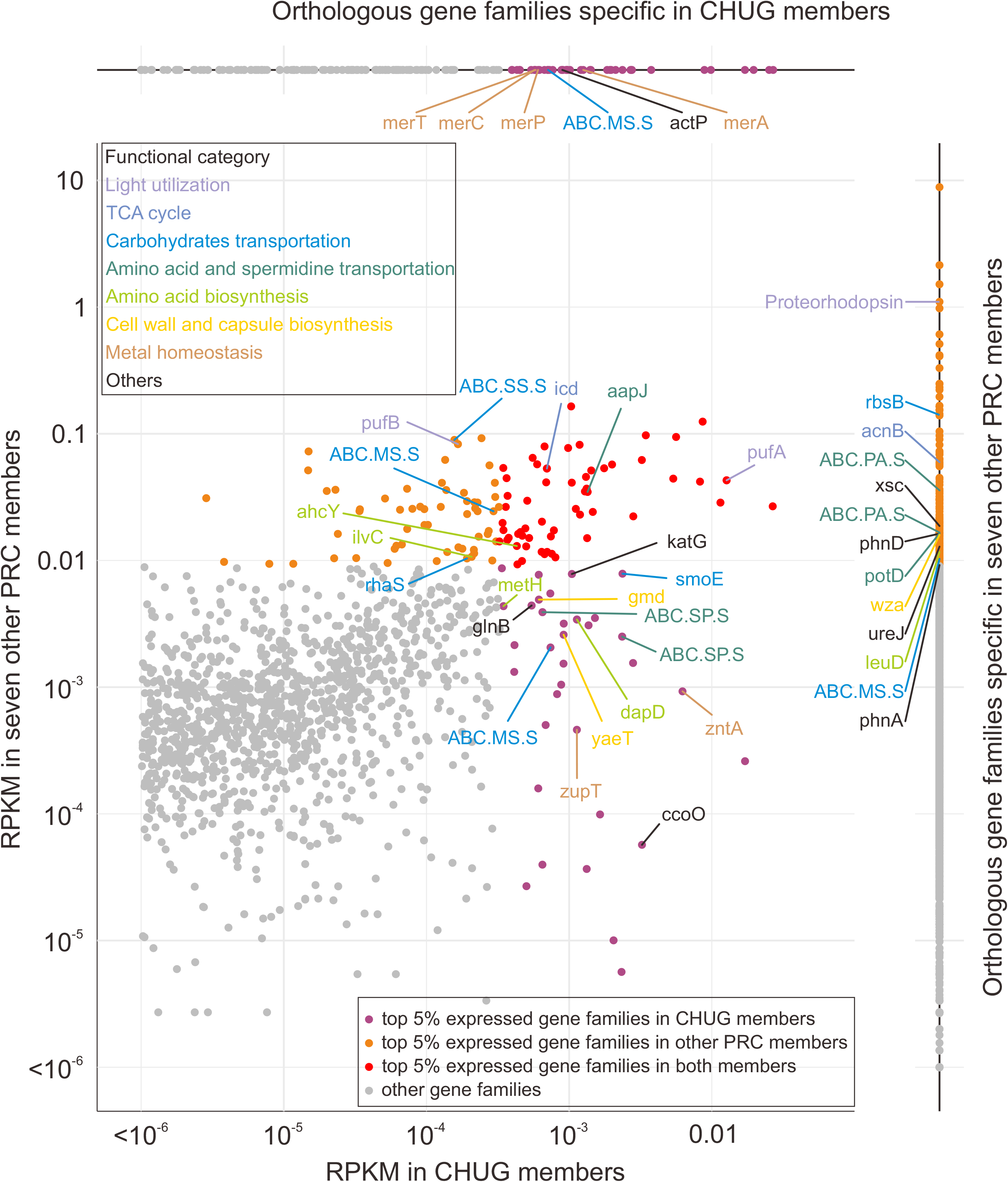
The average expression level of gene families in CHUG and seven other PRC members. Gene families with their gene expression level at top 5% found exclusively in CHUG members, exclusively in seven other PRC members, and shared by CHUG and other PRC members are shown in magenta, orange, and red dots, respectively. The remaining gene families are shown in gray dots. Gene families specific to CHUG and seven other PRC members are shown in the upper and right panel, respectively. Abbreviations: RPKM, Reads Per Kilobase per Million mapped reads; *aapJ*, general L-amino acid transport system; ABC.MS, multiple sugar transport system; ABC.PA, polar amino acid transport system; ABC.SP, spermidine/putrescine transport system; *acnB*, aconitate hydratase 2; *actP*, acetate permease; *ahcY*, adenosylhomocysteinase; ccoO, cytochrome *cbb_3_*-type oxidase; *dapD*, 2,3,4,5-tetrahydropyridine-2,6-dicarboxylate N-succinyltransferase; *glnB*, nitrogen regulatory protein P-II; *gmd*, GDPmannose 4,6-dehydratase; *icd*, isocitrate dehydrogenase; *ilvC*, ketol-acid reductoisomerase; *katG*, catalase-peroxidase; *leuD*, 3-isopropylmalate/(R)-2-methylmalate dehydratase; *merA*, mercuric reductase; *merC*, *merP* and *merT*, mercuric ion transport system; *metH*, 5-methyltetrahydrofolate--homocysteine methyltransferase; *phnA*, phosphonoacetate hydrolase; *phnD*, phosphonate transport system; *potD*, spermidine/putrescine transport system; *pufA* and *pufB*, light-harvesting complex 1; *rbsB*, ribose transport system; *rhaS*, rhamnose transport system; *ureJ*, urease; *wza*, polysaccharide biosynthesis/export protein; *xsc*, sulfoacetaldehyde acetyltransferase; *yaeT*, Outer membrane protein assembly factor; *zntA*, lead, cadmium, zinc and mercury transporting ATPase; *zupT*, zinc transporter.

### Genome reduction and vitamin B12 auxotrophy

Since CHUG has a well-supported sister group and outgroup (Fig. 1A), we reconstructed the gene gain and loss events that were associated with the origin of the CHUG cluster (Fig. 5A). The last common ancestor (LCA) of the CHUG cluster was estimated to have 2,320 genes, 2,134 orthologous gene families (1.09 gene copy per family), and a genome size of 2.35 Mb. There were 172 families (185 genes) gained and 406 families (425 genes) lost on the ancestral branch leading to the LCA of CHUG, while 28 and 52 families (30 and 79 genes) underwent copy number increase and decrease, respectively. Compared to its sister group and the outgroup, CHUG members lost 412 Kb (9.8%) on the ancestral branch leading to its LCA (filled triangle in Fig. 5A).

**Fig. 5.**
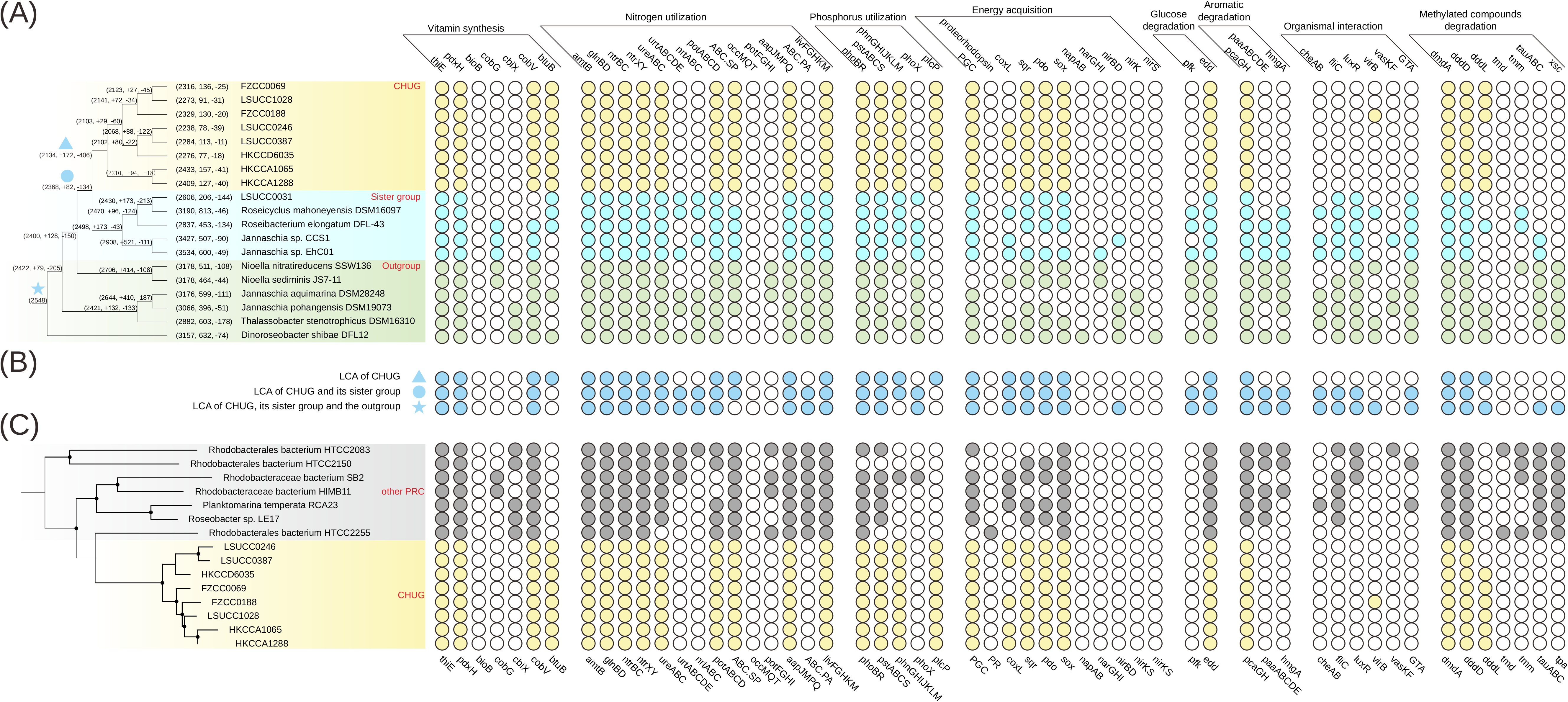
The phyletic pattern of select genes. The solid and open circles in the right panel represent the presence/absence of the genes, respectively. **(A)** The phyletic pattern in the CHUG, its sister group and its outgroup. The phylogenomic tree shown in the left panel is pruned from the full phylogenomic tree shown in Fig. 1A, and branch length is ignored for better visualization. The ancestral genome reconstruction was performed with BadiRate (32). Each ancestral and leaf node is associated with three numbers, representing the total number of orthologous gene families at this node, and the number of orthologous gene families gained and lost on the branch leading to this node. The LCA of CHUG, the LCA shared by CHUG and its sister group, and the LCA shared by CHUG, its sister group and the outgroup are marked with a filled triangle, a filled circle, and a filled star, respectively. **(B)** The estimated phyletic pattern of the above-mentioned three LCAs. **(C)** The gene presence and absence pattern in the CHUG and other seven PRC genomes. The dendrogram in the left panel is pruned from that shown in Fig. 1B. Abbreviations: *thiE*, thiamine-phosphate pyrophosphorylase; *pdxH*, pyridoxamine 5’-phosphate oxidase; *bioB*, biotin synthase; *cobG*, precorrin-3B synthase; *cbiX*, sirohydrochlorin cobaltochelatase; *cobV*, adenosylcobinamide-GDP ribazoletransferase; *btuB*, vitamin B12 transporter; *amtB*, ammonium transport system; nitrogen regulatory protein P-II (*glnBD*); *ntrBC*, nitrogen regulation two-component system; *ntrXY*, nitrogen regulation two-component system; *ureABC*, urease; *urtABCDE*, urea transport system; *nrtABC*, nitrate/nitrite transport system; *phoBR*, two-component phosphate regulatory system; *pstABCS*, phosphate transport system (high affinity); *phnGHIJKLM*, carbon-phosphorus (C-P) lyase; *phoX*, alkaline phosphatase; *plcP*, phospholipase C; PGC, photosynthesis gene cluster; *coxL*, carbon monoxide dehydrogenase (type I forming); *sqr*, sulfide quinone oxidoreductase; *pdo*, persulfide dioxygenase; *sox*, thiosulfate oxidizing SOX complex; *napAB*, nitrate reductase (periplasmic); *narGHI*, nitrate reductase (membrane-bound); *nirBD*, nitrite reductase; *nirK*, copper-containing NO-forming nitrite reductase; *nirS*, haem-containing NO-forming nitrite reductase; *pfk*, phosphofructokinase; *edd*, phosphogluconate dehydratase; *pcaGH*, protocatechuate 3,4-dioxygenase; *paaABCDE*, ring-1,2-phenylacetyl-CoA epoxidase; *hmgA*, homogentisate 1,2-dioxygenase; *cheAB*, chemotaxis family protein; *fliC*, flagellin; *luxR*, quorum-sensing system regulator; *virB*, type IV secretion system protein; va*sKF*, type VI secretion system protein; GTA, gene transfer agent; *dmdA*, DMSP demethylase; *dddD*, DMSP acyl-CoA transferase; *dddL*, dimethylpropiothetin dethiomethylase; *tmd*, trimethylamine dehydrogenase; *tmm*, trimethylamine monooxygenase; *tauABC*, taurine transport system; *xsc*, sulfoacetaldehyde acetyltransferase.

We further compared the metabolic potential between CHUG (Fig. 5A), the reconstructed ancestors (Fig. 5B), seven other PRC genomes (Fig. 5C), and other reference roseobacters (Table S2). Since the CHUG genomes experienced net DNA and gene losses, we explored whether metabolic auxotrophies (i.e., inability to synthesize a compound required for the growth) arose as a result of these losses. Among the sequenced CHUG members, the genome of the strain HKCCA1288 was closed, which improved our auxotrophy inference. CHUG genomes harbored the complete pathways for the synthesis of all 20 amino acids, many of which, such as the synthesis of lysine (*dapD*) and methionine (*metH* and *ahcY*), were under active expression in the wild (Fig. 4). They further encoded the key genes for thiamine (vitamin B_1_) synthesis (thiamine-phosphate pyrophosphorylase, *thiE*) and pyridoxine (vitamin B_6_) synthesis (pyridoxamine 5’-phosphate oxidase, *pdxH*). Nevertheless, the key gene for biotin (vitamin B_7_) synthesis (biotin synthase, *bioB*) was not found in CHUG nor in the sister group and the outgroup, suggesting that the biotin auxotrophy in CHUG was not part of their net gene losses.

Intriguingly, CHUG was auxotrophic for cobalamin (vitamin B_12_) biosynthesis, which can be synthesized by most roseobacters (2). This was validated using a growth assay, in which CHUG strain HKCCA1288 did not grow in the defined medium lacking vitamin B_12_ but grew well with the supplement of vitamin B_12_ (Fig. 6A). As a comparison, the model roseobacter *Ruegeria pomeroyi* DSS-3, which is equipped with the *cobG* route, grew equally well in the presence or absence of vitamin B_12_ (Fig. 6B). Mapping of the vitamin B_12_ *de novo* synthesis to the phylogeny (Fig. 1A) indicates that the loss of this capability was most likely associated with the genome reduction leading to the LCA of the CHUG lineage. On the other hand, no genome content changes were inferred related to vitamin B_12_ synthesis by the ancestral genome reconstruction (Fig. 5B). This controversy can be ascribed to the fact that the *de novo* synthesis of cobinamide has two non-homologous pathways (i.e., aerobic and anaerobic synthesis of cobinamide, the key precursor of vitamin B_12_, via key genes *cobG* and *cbiX*, respectively), and distinct pathways are maintained in the CHUG sister lineages (Fig. 1A). The ancestral genome reconstruction further inferred that the loss of vitamin B_12_ *de novo* synthesis capability was compensated with the coincidental gain of a putative vitamin B_12_ transporter (Fig. 5B), which was absent from all other PRC members capable of *de novo* vitamin B_12_ synthesis (Fig. 5C). Taken together, the loss of *de novo* synthesis capability and the gain of a putative transporter indicates that CHUG may have to acquire vitamin B_12_ or its precursor from the environment.

**Fig. 6.**
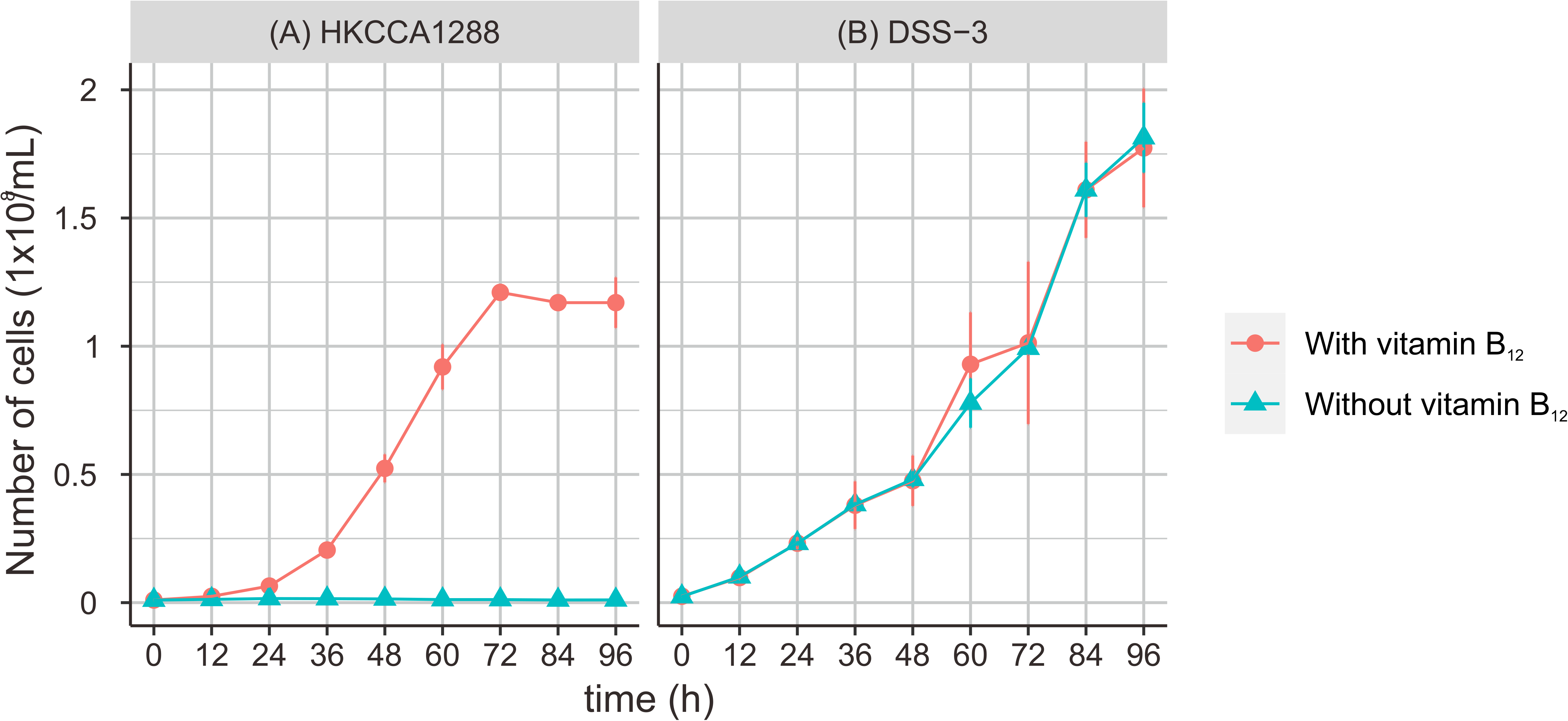
Growth assay of (A) CHUG strain HKCCA1288 and (B) model roseobacter *Ruegeria pomeroyi* DSS-3. Strains cultured on defined marine ammonium mineral salts (MAMS) medium with and without vitamin B_12_ were plotted in red and blue, respectively. Three triplicates were performed for each condition and error bars denote standard deviation.

### Metabolic potential for community interaction

Besides the loss of genes for *de novo* vitamin B_12_ synthesis, the CHUG members have also lost genes for chemotaxis (*cheAB*) and flagellar assembly (*fliC*). These genes were essential to mediate roseobacter-phytoplankton interactions (37), but may become dispensable when switching to a planktonic lifestyle (38). Consistent with this, the quorum-sensing (QS) system (*luxR*), type IV secretion system (*virB*), and type VI secretion system (*vasKF*) involved in organismal interactions were rarely found in the CHUG genomes (Fig. 5A). CHUG members also lost the gene cluster encoding gene transfer agent (GTA), which resembles small double-stranded DNA (dsDNA) bacteriophages that increase horizontal gene transfer (HGT) and metabolic flexibility at high population density (39).

### Metabolic potential for nutrient acquisition

Nitrogen (N) is a primarily limiting nutrient in surface oceans (40). Genes encoding the nitrogen regulatory protein P-II (*glnBD*) were highly expressed in the wild CHUG populations (Fig. 4). Genes encoding the high-affinity ammonium transporter (*amtB*) and nitrogen regulation two-component system (*ntrBC* and *ntrXY*) were found in the CHUG genomes. Genes encoding urease (*ureABC*) were also identified in CHUG members, though the urea transport system (*urtABCDE*) was not in any CHUG genomes. It is possible that urea was assimilated via passive diffusion across the cell membrane in CHUG as shown in other bacteria (41), or that urea was taken up by another promiscuous transporter. The genes encoding the transporter for nitrate/nitrite assimilation (*nrtABC*) were also missing in CHUG genomes. CHUG members retained the genes for the spermidine/putrescine transporter (*potABCD* and ABC.SP) (Table S2), and the latter was among the most highly expressed genes in the oceanic CHUG populations (Fig. 4). However, the CHUG members did not carry genes for other polyamine transport systems (*oocMQT* for octopine/nopaline and *potFGHI* for putrescine). The CHUG also retained and highly expressed *aapJMPQ* for the general L-amino acid transporter (Fig. 4), but lost genes encoding the polar amino acid transport system ABC.PA, which was prevalent in all other roseobacters studied here. CHUG further had a reduced number of genes (only one copy) encoding the branched-chain amino acid transport system (*livFGHKM*) compared to its sister group (at least three copies), the outgroup (at least three copies) and other PRC members (at least two copies; Table S2). Overall, fewer genes involved in the acquisition of amino acids were found in CHUG (Table S2), but they may remain efficient due to the high expression level of the retained genes.

Phosphorus (P) is often a co-limiting nutrient in surface oceans (40). To deal with P limitation, the CHUG members may be assisted by the essential regulatory and metabolic pathways known to be induced by P-limitation including the two-component regulatory system (*phoBR*), the high-affinity phosphate transporter (*pstABCS*) for phosphate acquisition and the C-P lyase (*phnGHIJKLM*) for phosphonate utilization. However, they have lost the *phoX* encoding an alkaline phosphatase for phosphodiester utilization (42) during the genome reduction process (Fig. 5A,B). A notable evolutionary innovation upon the emergence of the CHUG lineage was a gain of the gene encoding phospholipase C (*plcP*) (Fig. 5A,B), which was missing from all the seven other PRC members (Fig. 5C). The *plcP* is the key gene of the pathway for phospholipid substitution with non-phospholipids in response to P starvation, and was prevalently found in marine bacterioplankton (43).

### Metabolic potential for energy acquisition

Members of the CHUG cluster maintained some energy conservation strategies that are commonly found in other roseobacters. One example was the acquisition of light energy. The complete photosynthesis gene cluster underlying the aerobic anoxygenic photosynthesis (AAnP) were identified in all CHUG members, five of the seven other PRC genomes, and 21 of the 64 non-PRC genomes (Table S2). Other light energy acquisition mechanisms including genes encoding proteorhodopsin and xanthorhodopsin were only found in the PRC member HTCC2255 and in the two *Octadecabacter* genomes, respectively. Two marker genes (*pufAB*) of the photosynthesis gene cluster were among the most highly expressed genes in oceanic CHUG and other PRC members, and the proteorhodopsin in *Rhodobacterales* bacterium HTCC2255 was also highly expressed (Fig. 4). In total, the potential for light utilization was found in 14 of the 15 PRC members, but in only 23 of 64 non-PRC roseobacters (Table S2). The association of the light acquisition trait with the PRC members was significant, which remains when the biased phylogenetic distribution of this trait was under control as shown by the binaryPGLMM analysis (*p* < 0.05) (44). This result indicates that light utilization may facilitate their survival under nutrient-depleted pelagic environments (45, 46). However, it is not clear why the reduced CHUG genomes employ the photosynthesis gene cluster rather than a rhodopsin system for light acquisition, considering that the photosynthesis gene cluster consists of about 40 genes (46) whereas a rhodopsin system requires only the rhodopsin gene and an associated chromophore retinal gene (47). In fact, the possibility of an evolutionary replacement of photosynthesis gene cluster with proteorhodopsin remains open, because proteorhodopsin and photosynthesis gene cluster occur in two closely related ecotypes of DC5-80-3 (also called RCA), respectively (48), suggesting that the replacement of one phototrophic type with the other could happen rapidly.

Another example for energy conservation is the oxidation of reduced inorganic compounds. The CHUG carried genes for the oxidation of carbon monoxide (CO) and sulfide/thiosulfate as energy sources. Most roseobacters encode type II carbon monoxide dehydrogenase (*codh*), but only those with type I CODH may perform CO oxidation (49). Four of the eight CHUG genomes possessed type I CODH (*coxL*) and thus may oxidize CO (Fig. 5). This gene was further identified in three genomes from its sister group, three genomes from the outgroup, three PRC genomes and 18 other reference genomes (Table S2). All CHUG members possessed the sulfide:quinone oxidoreductase (*sqr*) for the oxidation of sulfide to zero valence sulfur (S^o^) (50), the persulfide dioxygenase (*pdo*) for the oxidation of S^o^ to sulfite which could spontaneous react with S^o^ to generate thiosulfate (50), and the complete *sox* pathway for the oxidation of thiosulfate to sulfate (51) (Fig. 5). The *sqr* and *pdo* were also found in four other PRC genomes and 32 of the 64 non-PRC genomes, while the *sox* pathway was found in all seven PRC genomes and 42 non-PRC genomes (Table S2). Unlike the capability of light utilization, no uneven distribution was identified for *coxL*, *sox*, *sqr* and *pdo* between PRC and non-PRC roseobacters (χ*2* test for *coxL* and binaryPGLMM analysis for the remaining genes; *p* > 0.05).

CHUG cannot perform nitrate/nitrite reduction for energy conservation due to the lack of genes involved in nitrate reduction to nitrite (nitrate reductase, periplasmic *napAB* or membrane-bound *narGHI*), nitrite reduction to ammonium (nitrite reductase, *nirBD*) or nitrite reduction to nitric oxide (NO-forming nitrite reductase, copper-containing *nirK* or haem-containing *nirS*) (2). The *narGHI* and *nirBD* were identified in some genomes affiliated with the sister group and the outgroup (Fig. 5A). These genes were also missing from other PRC genomes, but were found in some reference *Roseobacter* genomes (Table S2).

### Other important metabolic pathways relevant to Roseobacter ecology

Among the major pathways for glycolysis, all CHUG members maintained the key gene encoding phosphogluconate dehydratase (*edd*) for the Entner-Doudoroff (ED) pathway, but none of them contained the key gene for phosphofructokinase (*pfk*) in the Embden-Meyerhof-Parnas (EMP) pathway (Fig. 5). Both pathways were prevalent in the sister group and the outgroup. Ancestral genome content reconstruction inferred that the EMP pathway was lost at the LCA of the CHUG lineage (Fig. 5B) as a result of genome reduction. Interestingly, the seven other PRC genomes held an identical pattern to CHUG, in which the ED pathway was universally preserved but the EMP pathway was missing. Although generating less ATP and NADH, the ED pathway can provide NADPH and may accompany increased resistance to oxidative stress compared with the EMP pathway (52). This likely confers an important benefit to these pelagic roseobacters inhabiting the surface ocean where reactive oxygen species (ROS) production is intensive (53). The catabolic product of the ED pathway, pyruvate, can be further degraded through the tricarboxylic acid cycle (TCA) cycle, the genes of which were highly expressed in environmental CHUG members (Fig. 4).

Many roseobacters can degrade aromatic compounds through the aerobic ring-cleaving pathways (54). All CHUG members harbored the protocatechuate ring cleavage pathway (protocatechuate 3,4-dioxygenase, *pcaGH*), which is one of the most common pathways for the degradation of monoaromatic compounds among roseobacters (55). However, they did not carry *paaABCDE* encoding ring-1,2-phenylacetyl-CoA epoxidase (key enzyme for the phenylacetate ring cleavage pathway) or *hmgA* encoding homogentisate 1,2-dioxygenase (key enzyme for the homogenisate ring cleavage pathway) (54). As these two pathways were inferred to be present in the LCA shared by CHUG and its sister group (filled circle in Fig. 5A), we hypothesize that their absence from CHUG resulted from genome reduction. All the three ring cleavage pathways were common in the seven other PRC genomes (Table S2).

Methylated compounds are important substrates for roseobacters (56). Briefly, the CHUG members possessed the metabolic potential to utilize dimethylsulfoniopropionate (DMSP) via both demethylation (DMSP demethylase, *dmdA*) and cleavage (*dddD* or *dddL*) pathway. However, genes encoding trimethylamine dehydrogenase (*tmd*) and trimethylamine monooxygenase (*tmm*) involved in trimethylamine N-oxide (TMAO) and trimethylamine (TMA) degradation, respectively, were not identified in the CHUG genomes, nor in most genomes affiliated with their sister group and the outgroup. However, these genes were identified in some other PRC members. Genes involved in taurine transport (*tauABC*) and degradation (*xsc*) were not found in CHUG members, but they were present, and the latter was highly expressed, in seven other PRC members (Fig. 4).

## Discussion

### The CHUG population dynamics are uncoupled from phytoplankton abundance

Though the novel lineage CHUG and the previously known Pelagic *Roseobacter* Cluster (PRC) members all reach high global abundance and activity, the ecological factors driving their global distribution are different. DC5-80-3 and NAC11-7 abundances were previously shown to be positively correlated with phytoplankton blooms (1, 4, 57–60) and their abundance and activity were both found to be significantly correlated with Chl-a abundance here (Fig. 3D,E). In the PRC lineage CHAB-I-5, a few members carry signature genes mediating organismal interactions (e.g., type VI secretion system and quorum sensing) (12), and thus may also explore microenvironments such as phytoplankton and organic particles. In fact, CHAB-I-5 abundance and activity was also positively correlated with Chl-a across the global ocean samples (Fig. 3D,E), though such a correlation was not found in a previous study with a more limited sampling effort (11). In the case of CHUG, no significant correlation with Chl-a was identified (Fig. 3D,E). Indeed, when the *TARA Ocean* metagenomic sequencing reads at the nanoplankton-enriched size fraction (5-20 μm) were recruited, CHUG members exhibited a lower relative abundance than the other PRC representatives by approximately one order of magnitude (Fig. 3C). Together, these data support our hypothesis that members of the CHUG lineage evolved a free-living lifestyle decoupled from phytoplankton.

The possible contrasting roles of CHUG versus other pelagic roseobacters in relationship to phytoplankton were further supported by the absence of the *de novo* vitamin B_12_ synthesis in all CHUG members but its presence in all other PRC members. The auxotrophy for vitamin B_12_ was also validated for HKCCA1288 - for which we generated a complete genome sequence - in a growth assay (Fig. 6). The marine eukaryotic algae are predominantly vitamin B_12_ auxotrophs (61), whereas most roseobacters have the potential to synthesize vitamin B_12_ (2). This complementarity is one of the major mechanisms that facilitate mutualistic interactions between roseobacters and phytoplankton (2, 13–15), which also helps explain why roseobacters are often among the most abundant bacteria associated with marine eukaryotic phytoplankton (62–64). The loss of vitamin B_12_ synthesis in CHUG is unusual because members of the *Roseobacter* group are known as the dominant bacterial lineages associated with marine phytoplankton groups (65) and their evolutionary history was likely correlated with phytoplankton diversification (2, 66). They usually benefit from the fixed carbon or other excretions released by phytoplankton and, in return, produce secondary metabolites (e.g. vitamins, indole-3-acetic acid) to promote phytoplankton growth (15, 67, 68). These interactions likely occur in microzones immediately surrounding phytoplankton cells, which may create gene flow barriers and facilitate population differentiation of associated roseobacters (69). Therefore, the ecology and evolution of the *Roseobacter* group in the pelagic ocean are generally shaped by marine phytoplankton, making the possible separation from this ecological pattern in the CHUG lineage unique.

### Potential evolutionary forces driving genome reduction of the CHUG roseobacters

The most abundant marine bacterioplankton, such as the *Pelagibacterales* (also called the SAR11 clade) in the Alphaproteobacteria and the *Prochlorococcus* in Cyanobacteria, are often equipped with very small genomes (70). The evolutionary mechanisms driving their genome reduction have been discussed extensively. Among these, positive selection for metabolic efficiency (i.e., ‘genome streamlining’) has been theorized as the dominant force driving their genome reduction (70, 71). Although CHUG members possessed smaller genomes and lower GC content compared to the sister group and the outgroup, they did not show other features of genome streamlining, such as higher coding density, fewer paralogues, or a lower proportion of pseudogenes (70, 72). Therefore, the genome reduction process of CHUG members did not meet the canonical definition of ‘genome streamlining’.

Other important evidence against the genome streamlining explanation for CHUG genome reduction was from the genomic proxies for nutrient acquisition and saving strategies used by marine bacterioplankton. Among the selective factors that may drive bacterioplankton genome reduction in the pelagic ocean, N limitation is considered as the dominant factor (34, 70, 73, 74). Although the relative abundance of gene transcripts (but not the genes) in the wild CHUG populations was positively correlated with the nitrate concentration (Fig. 3E), which provides marginal evidence for a role of N limitation, other key evidence was missing. For example, we did not observe a reduced use of N in the amino acid sequences (approximated by N-ARSC) in CHUG compared to the sister group and the outgroup. Similar observation was used as evidence against the hypothesis that N limitation is a strong driver of genome streamlining in other marine bacterioplankton lineages (75, 76). A second potential ecological factor driving genome streamlining is P limitation (77), though this theory has been debated (78). Genome reduction likely leads to a sizable decrease in cellular P requirement and thus may confer a competitive advantage in the P-limited marine environments (79). Although a few important genes for P acquisition (*pst* for high-affinity phosphate transporter and *phn* for C-P lyase) were retained during the CHUG genome reduction and a gene encoding phospholipase C (*plcP*) responsible for cell membrane phospholipid substitution for non-phosphorus lipids (43) was even acquired, the key P scavenging gene encoding PhoX alkaline phosphatase was lost (Fig. 5). Therefore, available evidence for either N or P limitation as a driver of CHUG genome reduction was self-contradictory.

Because evidence for genome streamlining was weak in this lineage, we examined neutral evolutionary forces as potential explanations for CHUG genome reduction. In fact, neutral mechanisms have recently been considered to play an important role in driving genome reduction of marine bacterioplankton lineages (80–82). Most of the prior studies focused on *Prochlorococcus* (see references cited in the following paragraphs). While some extended their discussions to *Pelagibacterales* (81, 83), knowledge on the evolutionary mechanisms driving genome reduction of most other marine bacterioplankton lineages is rather limited.

One potentially important neutral driver is random genetic drift due to a reduction of effective population size (*N_e_*). A previous study showed that the major genome reduction event coincided with an accelerated rate of accumulating deleterious mutations in the early evolution of *Prochlorococcus*, providing important evidence that genetic drift was likely the primary mechanism of genome reduction in this lineage (81). Specifically, the power of genetic drift (i.e., the inverse of *N_e_*) of an ancestral lineage (e.g., the ancestral branch underlying the ancient genomic events) can be approximated by the ratio of the radical nonsynonymous nucleotide substitutions per radical nonsynonymous site (*d_R_*) to the conservative nonsynonymous nucleotide substitutions per conservative nonsynonymous site (*d_C_*) (81). Because the replacement by a physicochemically dissimilar amino acid (or radical change) is likely to be more deleterious than the replacement by a similar amino acid (or conservative change) (84, 85), the elevated *d_R_*/*d_C_* ratio is evidence for genetic drift acting to accumulate the deleterious type of mutations (i.e., the radical changes) in excess. In terms of the CHUG, the *d_R_*/*d_C_* ratio was not significantly elevated compared to its sister group (Fig. S5A) under two independent methods for biochemical classification of the 20 amino acids (Fig. S5B,C), suggesting that the deleterious type of mutations was not accumulated in excess at the ancestral branch leading to the LCA of the CHUG lineage (filled triangle in Fig. 5A). Since this ancestral branch corresponds to the time when major genome reduction occurred for CHUG, we can conclude that genetic drift was unlikely to be an important driver of CHUG genome reduction.

A second potentially important neutral driver of prokaryotic genome reduction is increased mutation rate, which was also proposed to explain *Prochlorococcus* genome reduction (86). Mathematical modeling predicts that not all auxiliary genes can be maintained by purifying selection when mutation rate is increased, and that an increase of 10 fold in mutation rate may lead to a 30% decrease in genome size (87). More recently, this hypothesis was supported with empirical data from comparative genomics analyses (82), though whether increased mutation rate is a truly important driver of prokaryotic genome reduction is debated (88). Given the potentially important role of increased mutation rate in driving prokaryotic genome reduction, determining the unbiased spontaneous mutation rate of the CHUG and the sister lineage using the mutation accumulation experiment followed by whole genome sequencing of the mutant lines becomes an urgent research need.

One more potentially important but rarely discussed neutral force leading to genome reduction is the loss of the genes that were important in the initial habitat but became dispensable after the bacteria switched to a new environment. This neutral loss mechanism, termed relaxation of purifying selection, may also have contributed to genome reduction in *Prochlorococcus* (89). Importantly, the loss of dispensable genes under this mechanism is not related to the change of *N_e_* but results instead from a change of habitat or lifestyle. Unlike other pelagic roseobacter members, CHUG members do not exhibit a correlative pattern between their global distributions and Chl-a (Fig. 3D,E), which can be used as a proxy for phytoplankton abundances (90). This is supported by evidence at the molecular and physiological level, in which the *de novo* synthesis of vitamin B_12_, a fundamental metabolite roseobacters produce and supply to phytoplankton, was missing from all CHUG members but present in all other pelagic roseobacters (Fig. 1A, Fig. 6). Once the capability of *de novo* vitamin B_12_ synthesis was lost, the CHUG ancestor may have lost its ability to establish symbiosis with phytoplankton and subsequently undergone a major shift of its planktonic lifestyle, namely from phytoplankton-associated to free-living. Given that phytoplankton cell surfaces can be more densely populated compared to the bulk seawater (65), genes contributing to roseobacter-phytoplankton symbiosis (e.g., motility and chemotaxis), depending on population density and involved in interactions with other bacteria (e.g., quorum sensing, gene transfer agent), may have become dispensable during this transition (38). Indeed, the loss of these signature genes contributed to the genome reduction of CHUG (Fig. 5). We therefore propose that the relaxation of purifying selection may be one of the primary evolutionary forces leading to the major genome reduction of CHUG.

## Data availability

Genomic sequences of the eight CHUG genomes are available at the NCBI GenBank database under the accession number PRJNA574877.

## Code availability

The custom scripts used in this study are available in the online repository (https://github.com/luolab-cuhk/CHUG-genome-reduction-project).

## Acknowledgments

This research was funded by the National Science Foundation of China (41776129), the Hong Kong Research Grants Council General Research Fund (14163917), the Hong Kong Research Grants Council Area of Excellence Scheme (AoE/M-403/16), and the Direct Grant of CUHK (4053257 & 3132809). The research was also supported by a Louisiana Board of Regents grant (LEQSF(2014-17)-RD-A-06) and a Simons Early Career Investigator in Marine Microbial Ecology and Evolution Award to JCT.

## Conflict of Interest

The authors declare no competing interests concerning the submitted work.

## Notes

### Competing Interest Statement

The authors have declared no competing interest.

## References

1. Buchan A, González JM, Moran MA. Overview of the marine *Roseobacter* lineage. Appl Environ Microbiol 2005; 71(10):5665–77.

2. Luo H, Moran MA. Evolutionary ecology of the marine *Roseobacter* clade. Microbiol Mol Biol Rev 2014; 78(4):573–87.

3. Moran MA, Belas R, Schell MA, González JM, Sun F, Sun S et al. Ecological genomics of marine Roseobacters. Appl Environ Microbiol 2007; 73(14):4559–69.

4. Giebel H-A, Kalhoefer D, Lemke A, Thole S, Gahl-Janssen R, Simon M et al. Distribution of *Roseobacter* RCA and SAR11 lineages in the North Sea and characteristics of an abundant RCA isolate. ISME J 2011; 5(1):8–19.

5. Wemheuer B, Wemheuer F, Hollensteiner J, Meyer F-D, Voget S, Daniel R. The green impact: bacterioplankton response toward a phytoplankton spring bloom in the southern North Sea assessed by comparative metagenomic and metatranscriptomic approaches. Front Microbiol 2015; 6:805.

6. Pujalte MJ, Lucena T, Ruvira MA, Arahal DR, Macián MC. The Family *Rhodobacteraceae*. In: Rosenberg E, DeLong EF, Lory S, Stackebrandt E, Thompson F, editors. The Prokaryotes. Berlin, Heidelberg: Springer Berlin Heidelberg; 2014. p. 439–512.

7. Buchan A, Hadden M, Suzuki MT. Development and application of quantitative-PCR tools for subgroups of the *Roseobacter* clade. Appl Environ Microbiol 2009; 75(23):7542–7.

8. Luo H, Swan BK, Stepanauskas R, Hughes AL, Moran MA. Comparing effective population sizes of dominant marine alphaproteobacteria lineages. Environ Microbiol Rep 2014; 6(2):167–72.

9. Giebel H-A, Kalhoefer D, Gahl-Janssen R, Choo Y-J, Lee K, Cho J-C et al. *Planktomarina temperata* gen. nov., sp. nov., belonging to the globally distributed RCA cluster of the marine *Roseobacter* clade, isolated from the German Wadden Sea. Int J Syst Evol Microbiol 2013; 63(Pt 11):4207–17.

10. Voget S, Wemheuer B, Brinkhoff T, Vollmers J, Dietrich S, Giebel H-A et al. Adaptation of an abundant *Roseobacter* RCA organism to pelagic systems revealed by genomic and transcriptomic analyses. ISME J 2015; 9(2):371–84.

11. Billerbeck S, Wemheuer B, Voget S, Poehlein A, Giebel H-A, Brinkhoff T et al. Biogeography and environmental genomics of the *Roseobacter*-affiliated pelagic CHAB-I-5 lineage. Nat Microbiol 2016; 1(7):16063.

12. Zhang Y, Sun Y, Jiao N, Stepanauskas R, Luo H. Ecological genomics of the uncultivated marine *Roseobacter* lineage CHAB-I-5. Appl Environ Microbiol 2016; 82(7):2100–11.

13. Wagner-Döbler I, Ballhausen B, Berger M, Brinkhoff T, Buchholz I, Bunk B et al. The complete genome sequence of the algal symbiont *Dinoroseobacter shibae*: a hitchhiker’s guide to life in the sea. ISME J 2010; 4(1):61–77.

14. Durham BP, Sharma S, Luo H, Smith CB, Amin SA, Bender SJ et al. Cryptic carbon and sulfur cycling between surface ocean plankton. Proc Natl Acad Sci U S A 2015; 112(2):453–7.

15. Cooper MB, Kazamia E, Helliwell KE, Kudahl UJ, Sayer A, Wheeler GL et al. Cross-exchange of B-vitamins underpins a mutualistic interaction between *Ostreococcus tauri* and *Dinoroseobacter shibae*. ISME J 2018.

16. Bankevich A, Nurk S, Antipov D, Gurevich AA, Dvorkin M, Kulikov AS et al. SPAdes: a new genome assembly algorithm and its applications to single-cell sequencing. J Comput Biol 2012; 19(5):455–77.

17. Seemann T. Prokka: rapid prokaryotic genome annotation. Bioinformatics 2014; 30(14):2068–9.

18. Jain C, Rodriguez-R LM, Phillippy AM, Konstantinidis KT, Aluru S. High throughput ANI analysis of 90K prokaryotic genomes reveals clear species boundaries. Nat Commun 2018; 9(1):5114.

19. Parks DH, Imelfort M, Skennerton CT, Hugenholtz P, Tyson GW. CheckM: assessing the quality of microbial genomes recovered from isolates, single cells, and metagenomes. Genome Res 2015; 25(7):1043–55.

20. Parks DH, Rinke C, Chuvochina M, Chaumeil P-A, Woodcroft BJ, Evans PN et al. Recovery of nearly 8,000 metagenome-assembled genomes substantially expands the tree of life. Nat Microbiol 2017; 2(11):1533–42.

21. Chu X, Li S, Wang S, Luo D, Luo H. Gene loss through pseudogenization contributes to the ecological diversification of a generalist *Roseobacter* lineage. ISME J 2020.

22. Revell LJ. phytools: an R package for phylogenetic comparative biology (and other things). Methods in Ecology and Evolution 2012; 3(2):217–23.

23. Sunagawa S, Coelho LP, Chaffron S, Kultima JR, Labadie K, Salazar G et al. Ocean plankton. Structure and function of the global ocean microbiome. Science 2015; 348(6237):1261359.

24. Salazar G, Paoli L, Alberti A, Huerta-Cepas J, Ruscheweyh H-J, Cuenca M et al. Gene expression changes and community turnover differentially shape the global ocean metatranscriptome. Cell 2019; 179(5):1068–1083.e21.

25. Vargas C de, Audic S, Henry N, Decelle J, Mahé F, Logares R et al. Eukaryotic plankton diversity in the sunlit ocean. Science 2015; 348(6237):1261605.

26. Langmead B, Salzberg SL. Fast gapped-read alignment with Bowtie 2. Nat Methods 2012; 9(4):357–9.

27. Altschul SF, Gish W, Miller W, Myers EW, Lipman DJ. Basic local alignment search tool. J Mol Biol 1990; 215(3):403–10.

28. Harrell Jr FE. Package ‘Hmisc’. CRAN2018 2019; 2019:235–6.

29. Parks DH, Chuvochina M, Waite DW, Rinke C, Skarshewski A, Chaumeil P-A et al. A standardized bacterial taxonomy based on genome phylogeny substantially revises the tree of life. Nat Biotechnol 2018; 36(10):996–1004.

30. Simon M, Scheuner C, Meier-Kolthoff JP, Brinkhoff T, Wagner-Döbler I, Ulbrich M et al. Phylogenomics of *Rhodobacteraceae* reveals evolutionary adaptation to marine and non-marine habitats. ISME J 2017; 11(6):1483–99.

31. Emms DM, Kelly S. OrthoFinder: phylogenetic orthology inference for comparative genomics. Genome Biol 2019; 20(1):238.

32. Librado P, Vieira FG, Rozas J. BadiRate: estimating family turnover rates by likelihood-based methods. Bioinformatics 2012; 28(2):279–81.

33. Lekunberri I, Gasol JM, Acinas SG, Gómez-Consarnau L, Crespo BG, Casamayor EO et al. The phylogenetic and ecological context of cultured and whole genome-sequenced planktonic bacteria from the coastal NW Mediterranean Sea. Syst Appl Microbiol 2014; 37(3):216–28.

34. Luo H, Swan BK, Stepanauskas R, Hughes AL, Moran MA. Evolutionary analysis of a streamlined lineage of surface ocean *Roseobacters*. ISME J 2014; 8(7):1428–39.

35. Coello-Camba A, Diaz-Rua R, Duarte CM, Irigoien X, Pearman JK, Alam IS et al. Picocyanobacteria community and cyanophage infection responses to nutrient enrichment in a mesocosms experiment in oligotrophic waters. Front. Microbiol. 2020; 11.

36. Kim Y, Jeon J, Kwak MS, Kim GH, Koh I, Rho M. Photosynthetic functions of *Synechococcus* in the ocean microbiomes of diverse salinity and seasons. PLoS ONE 2018; 13(1):e0190266.

37. Geng H, Belas R. Molecular mechanisms underlying *Roseobacter*phytoplankton symbioses. Curr Opin Biotechnol 2010; 21(3):332–8.

38. Luo H, Moran MA. How do divergent ecological strategies emerge among marine bacterioplankton lineages? Trends Microbiol 2015; 23(9):577–84.

39. Biers EJ, Wang K, Pennington C, Belas R, Chen F, Moran MA. Occurrence and expression of gene transfer agent genes in marine bacterioplankton. Appl Environ Microbiol 2008; 74(10):2933–9.

40. Moore CM, Mills MM, Arrigo KR, Berman-Frank I, Bopp L, Boyd PW et al. Processes and patterns of oceanic nutrient limitation. Nature Geosci 2013; 6(9):701–10.

41. Veaudor T, Cassier-Chauvat C, Chauvat F. Genomics of urea transport and catabolism in Cyanobacteria: biotechnological implications. Front Microbiol 2019; 10:2052.

42. Luo H, Benner R, Long RA, Hu J. Subcellular localization of marine bacterial alkaline phosphatases. Proc Natl Acad Sci U S A 2009; 106(50):21219–23.

43. Sebastián M, Smith AF, González JM, Fredricks HF, van Mooy B, Koblížek M et al. Lipid remodelling is a widespread strategy in marine heterotrophic bacteria upon phosphorus deficiency. ISME J 2016; 10(4):968–78.

44. Paradis E, Schliep K. ape 5.0: an environment for modern phylogenetics and evolutionary analyses in R. Bioinformatics 2019; 35(3):526–8.

45. Yooseph S, Nealson KH, Rusch DB, McCrow JP, Dupont CL, Kim M et al. Genomic and functional adaptation in surface ocean planktonic prokaryotes. Nature 2010; 468(7320):60–6.

46. Brinkmann H, Göker M, Koblížek M, Wagner-Döbler I, Petersen J. Horizontal operon transfer, plasmids, and the evolution of photosynthesis in *Rhodobacteraceae*. ISME J 2018; 12(8):1994–2010.

47. Pinhassi J, DeLong EF, Béjà O, González JM, Pedrós-Alió C. Marine Bacterial and Archaeal Ion-Pumping Rhodopsins: Genetic Diversity, Physiology, and Ecology. Microbiol Mol Biol Rev 2016; 80(4):929–54.

48. Sun Y, Zhang Y, Hollibaugh JT, Luo H. Ecotype diversification of an abundant *Roseobacter* lineage. Environ Microbiol 2017; 19(4):1625–38.

49. Cunliffe M. Correlating carbon monoxide oxidation with cox genes in the abundant Marine *Roseobacter* Clade. ISME J 2011; 5(4):685–91.

50. Xin Y, Liu H, Cui F, Liu H, Xun L. Recombinant *Escherichia coli* with sulfide:quinone oxidoreductase and persulfide dioxygenase rapidly oxidises sulfide to sulfite and thiosulfate via a new pathway. Environ Microbiol 2016; 18(12):5123–36.

51. Friedrich CG, Quentmeier A, Bardischewsky F, Rother D, Kraft R, Kostka S et al. Novel genes coding for lithotrophic sulfur oxidation of *Paracoccus pantotrophus* GB17. J. Bacteriol. 2000; 182(17):4677–87.

52. Klingner A, Bartsch A, Dogs M, Wagner-Döbler I, Jahn D, Simon M et al. Large-Scale 13C flux profiling reveals conservation of the Entner-Doudoroff pathway as a glycolytic strategy among marine bacteria that use glucose. Appl Environ Microbiol 2015; 81(7):2408–22.

53. Diaz JM, Hansel CM, Voelker BM, Mendes CM, Andeer PF, Zhang T. Widespread production of extracellular superoxide by heterotrophic bacteria. Science 2013; 340(6137):1223–6.

54. Gulvik CA, Buchan A. Simultaneous catabolism of plant-derived aromatic compounds results in enhanced growth for members of the *Roseobacter* lineage. Appl Environ Microbiol 2013; 79(12):3716–23.

55. Alejandro-Marín CM, Bosch R, Nogales B. Comparative genomics of the protocatechuate branch of the β-ketoadipate pathway in the *Roseobacter* lineage. Marine Genomics 2014; 17:25–33.

56. Chen Y. Comparative genomics of methylated amine utilization by marine *Roseobacter* clade bacteria and development of functional gene markers (tmm, gmaS). Environ Microbiol 2012; 14(9):2308–22.

57. Wagner-Döbler I, Biebl H. Environmental biology of the marine *Roseobacter* lineage. Annu Rev Microbiol 2006; 60:255–80.

58. West NJ, Obernosterer I, Zemb O, Lebaron P. Major differences of bacterial diversity and activity inside and outside of a natural iron-fertilized phytoplankton bloom in the Southern Ocean. Environ Microbiol 2008; 10(3):738–56.

59. Rich VI, Pham VD, Eppley J, Shi Y, DeLong EF. Time-series analyses of Monterey Bay coastal microbial picoplankton using a ‘genome proxy’ microarray. Environ Microbiol 2011; 13(1):116–34.

60. Landa M, Blain S, Christaki U, Monchy S, Obernosterer I. Shifts in bacterial community composition associated with increased carbon cycling in a mosaic of phytoplankton blooms. ISME J 2016; 10(1):39–50.

61. Helliwell KE. The roles of B vitamins in phytoplankton nutrition: new perspectives and prospects. New Phytol 2017; 216(1):62–8.

62. González JM, Simó R, Massana R, Covert JS, Casamayor EO, Pedrós-Alió C et al. Bacterial community structure associated with a dimethylsulfoniopropionate-producing North Atlantic algal bloom. Appl Environ Microbiol 2000; 66(10):4237–46.

63. Amin SA, Parker MS, Armbrust EV. Interactions between diatoms and bacteria. Microbiol Mol Biol Rev 2012; 76(3):667–84.

64. Li S, Chen M, Chen Y, Tong J, Wang L, Xu Y et al. Epibiotic bacterial community composition in red-tide dinoflagellate *Akashiwo sanguinea* culture under various growth conditions. FEMS Microbiol Ecol 2019; 95(5).

65. Seymour JR, Amin SA, Raina J-B, Stocker R. Zooming in on the phycosphere: the ecological interface for phytoplankton-bacteria relationships. Nat Microbiol 2017; 2:17065.

66. Luo H, Csuros M, Hughes AL, Moran MA. Evolution of divergent life history strategies in marine alphaproteobacteria. MBio 2013; 4(4).

67. Durham BP, Dearth SP, Sharma S, Amin SA, Smith CB, Campagna SR et al. Recognition cascade and metabolite transfer in a marine bacteria-phytoplankton model system. Environ Microbiol 2017:3500–13.

68. Shibl AA, Isaac A, Ochsenkühn MA, Cárdenas A, Fei C, Behringer G et al. Diatom modulation of select bacteria through use of two unique secondary metabolites. Proc Natl Acad Sci U S A 2020; 117(44):27445–55.

69. Wang X, Zhang Y, Ren M, Xia T, Chu X, Liu C et al. Cryptic speciation of a pelagic *Roseobacter* population varying at a few thousand nucleotide sites. ISME J 2020.

70. Giovannoni SJ, Cameron Thrash J, Temperton B. Implications of streamlining theory for microbial ecology. ISME J 2014; 8(8):1553–65.

71. Giovannoni SJ, Tripp HJ, Givan S, Podar M, Vergin KL, Baptista D et al. Genome streamlining in a cosmopolitan oceanic bacterium. Science 2005; 309(5738):1242–5.

72. Swan BK, Tupper B, Sczyrba A, Lauro FM, Martinez-Garcia M, González JM et al. Prevalent genome streamlining and latitudinal divergence of planktonic bacteria in the surface ocean. Proc Natl Acad Sci U S A 2013; 110(28):11463–8.

73. Luo H, Thompson LR, Stingl U, Hughes AL. Selection maintains low genomic GC content in marine SAR11 lineages. Mol Biol Evol 2015; 32(10):2738–48.

74. Mende DR, Bryant JA, Aylward FO, Eppley JM, Nielsen T, Karl DM et al. Environmental drivers of a microbial genomic transition zone in the ocean’s interior. Nat Microbiol 2017; 2(10):1367–73.

75. Grzymski JJ, Dussaq AM. The significance of nitrogen cost minimization in proteomes of marine microorganisms. ISME J 2012; 6(1):71–80.

76. Lee MD, Ahlgren NA, Kling JD, Walworth NG, Rocap G, Saito MA et al. Marine *Synechococcus* isolates representing globally abundant genomic lineages demonstrate a unique evolutionary path of genome reduction without a decrease in GC content. Environ Microbiol 2019; 21(5):1677–86.

77. Hessen DO, Jeyasingh PD, Neiman M, Weider LJ. Genome streamlining and the elemental costs of growth. Trends Ecol Evol (Amst) 2010; 25(2):75–80.

78. Vieira-Silva S, Touchon M, Rocha EPC. No evidence for elemental-based streamlining of prokaryotic genomes. Trends Ecol Evol (Amst) 2010; 25(6):319–20; author reply 320-1.

79. Thingstad T, Rassoulzadegan F. Conceptual models for the biogeochemical role of the photic zone microbial food web, with particular reference to the Mediterranean Sea. Progress in Oceanography 1999; 44(1-3):271–86.

80. Batut B, Knibbe C, Marais G, Daubin V. Reductive genome evolution at both ends of the bacterial population size spectrum. Nat Rev Microbiol 2014; 12(12):841–50.

81. Luo H, Huang Y, Stepanauskas R, Tang J. Excess of non-conservative amino acid changes in marine bacterioplankton lineages with reduced genomes. Nat Microbiol 2017; 2:17091.

82. Bourguignon T, Kinjo Y, Villa-Martín P, Coleman NV, Tang Q, Arab DA et al. Increased mutation rate is linked to genome reduction in prokaryotes. Curr Biol 2020; 30(19):3848–3855.e4.

83. Viklund J, Ettema TJG, Andersson SGE. Independent genome reduction and phylogenetic reclassification of the oceanic SAR11 clade. Mol Biol Evol 2012; 29(2):599–615.

84. Zuckerkandl E, Pauling L, Bryson V, Vogel HJ. Evolving genes and proteins. Science 1965:68–71.

85. Dayhoff MO. Atlas of protein sequence and structure. National Biomedical Research Foundation; 1972.

86. Dufresne A, Garczarek L, Partensky F. Accelerated evolution associated with genome reduction in a free-living prokaryote. Genome Biol 2005; 6(2):R14.

87. Marais GAB, Calteau A, Tenaillon O. Mutation rate and genome reduction in endosymbiotic and free-living bacteria. Genetica 2008; 134(2):205–10.

88. Gu J, Wang X, Ma X, Sun Y, Xiao X, Luo H. Unexpectedly high mutation rate of a deep-sea hyperthermophilic anaerobic archaeon. ISME J 2021; In press.

89. Luo H, Friedman R, Tang J, Hughes AL. Genome reduction by deletion of paralogs in the marine cyanobacterium *Prochlorococcus*. Mol Biol Evol 2011; 28(10):2751–60.

90. Roesler C, Uitz J, Claustre H, Boss E, Xing X, Organelli E et al. Recommendations for obtaining unbiased chlorophyll estimates from in situ chlorophyll fluorometers: A global analysis of WET Labs ECO sensors. Limnol. Oceanogr. Methods 2017; 15(6):572–85.

91. Nguyen L-T, Schmidt HA, Haeseler A von, Minh BQ. IQ-TREE: a fast and effective stochastic algorithm for estimating maximum-likelihood phylogenies. Mol Biol Evol 2015; 32(1):268–74.

