## Supplementary information for "Mechanisms Driving Genome Reduction of a Novel *Roseobacter* Lineage Showing Vitamin B_12_ Auxotrophy"

**This PDF file includes:**

Text 1. Supplementary methods

Text 2. Supplementary results

Figures S1 to S5

References

Text 1. Supplementary methods

1.1 Sampling, bacterial cultivation and genome sequencing

1.2 Genome assembly and annotation

1.3 Phylogenomic analysis

1.4 Phylogenetic analysis based on 16S rRNA genes

1.5 Genome content analysis

1.6 Statistical analysis for genomic features

1.7 Recruitment analysis using public metagenomic sequencing data

1.8 $d_R/d_C$ ratio calculation

1.9 Vitamin $B_{12}$ assay

Text 2. Supplementary results

2.1 Biased codon and amino acid usage

**Text 1. Supplementary methods**

*1.1 Sampling, bacterial cultivation and genome sequencing*

HKCC strains were isolated from ambient seawater samples from both the coral *Platygyra acuta* ecosystem (lat. N22.52°, long. E114.3173°) and the brown alga *Sargassum hemiphyllum* ecosystem that were collected in 2017 by scuba diving near Hong Kong, China. These samples were stored in the 50 mL microcentrifuge tube at 4°C. Marine basal medium (MBM) was modified with 1 mM of dimethylsulfoniopropionate (DMSP) as the sole carbon source. A 100-fold serial dilution of seawater was prepared, and a 100 μL aliquot of the diluted sample was spread on the MBM-DMSP medium. The isolation plates were incubated at room temperature. Colonies were selected after one week and repeatedly re-streaked on the 2216E marine agar (BD Difco, USA) to collect the biomass for DNA extraction. Genomic DNA was extracted using EZ.N.A. Bacterial DNA Kit, and was sent to Qingdao Huada Gene Biotechnology Co., Ltd for library preparation and genome sequencing with BGISEQ500 (PE100) following the standard protocol (1). In addition, biomass of HKCCA1288 was sent to Qingdao Huada Gene Biotechnology Co., Ltd for library preparation and genome sequencing with PacBio Sequel platform following a previous study (2) to obtain a complete and closed genome.

FZCC strains were isolated from the coastal water of Pingtan Island (lat. N25.43°, long. E119.78°) in May 2017 using high-throughput dilution-to-extinction cultivation (HTC) method (3–5). The seawater sample was diluted with seawater-based media (4, 5) and dispensed into 24-well polystyrene microplates (Corning Incorporated) with a final inoculation density of 3 cells/well. The plates were incubated at 20˚C in the dark for four weeks and counted with flow cytometry after staining with SYBR Green I. Wells with at least $10^5$ cells/mL were defined as positive cultures. Cells (500 μl) of positive cultures were collected by centrifugation (10,000 rpm, 30 min) and identified by 16S rRNA gene sequence identity as described previously (5). FZCC strains used for genome sequencing were grown in polycarbonate flasks (50 mL culture volume) to a cell density of >$10^6$ cells/mL and collected by centrifugation. Genomic DNA was extracted using DNeasy Blood & Tissue Kit (Qiagen, Valencia, CA, USA), and was sent for genome sequencing using Illumina HiSeq 2500 (PE150) at Hubbard Center for Genome Studies, University of New Hampshire.

LSUCC strains were isolated as previously described (6, 7) with the exception of LSUCC1028, in which we used a modified JW1 medium (6) containing no carbon sources except for fulvic acids (Santa Cruz Biotechnology, Inc, TX, USA; CAS 479-66-3) at 1mg/L ("JW1FA"). LSUCC1028 was isolated from surface water collected from Terrebonne Bay, LA (lat. N29.1872°, long. W90.62822°) in July 2015. Briefly, the collected water was diluted in JW1FA and inoculated into a 96-well Teflon plate at a density of 2 cells/well. The plate was incubated in the dark at 24˚C for two weeks and counted with flow cytometry. Each well was transferred into a new plate containing fresh media and counted after one week. This was repeated once more for a total of three plates. Any wells that remained positive ($10^4$ cells/mL) after three consecutive plates were subjected to three rounds of serial dilution to ensure culture purity. Cultures were then grown in polycarbonate flasks (50 mL culture volume) to a cell density of $10^6$ cells/mL and identified by16S rRNA gene sequence identity as described described (6, 7). Genomic DNA was extracted from LSUCC0031 via the DNeasy kit (Qiagen), from LSUCC0246 and LSUCC0387 via phenol-chloroform, and from LSUCC1028 via the PowerWater kit (Mo Bio). Genomic DNA was sent for genome sequencing using Illumina HiSeq 2500 (PE150) at Hubbard Center for Genome Studies, University of New Hampshire.

#### *1.2 Genome assembly and annotation*

The Illumina sequencing raw reads were quality trimmed with Trimmomatic v0.36 (8) with options ‘SLIDINGWINDOW:4:15 MAXINFO:40:0.9 MINLEN:40’ and assembled using SPAdes v3.10.1 (9) with ‘-careful’ options. Only contigs with length >2,000 bp and sequencing depth >5x were retained.

The quality of sequenced PacBio reads for HKCCA1288 was checked using FastQC v.0.11.4 (10). This strain was assembled by combining the Illumina short reads and the PacBio long reads using Unicycler v0.4.6 (11) with default parameters.

Genome completeness, contamination, and strain heterogeneity (Table S1) were calculated using CheckM v1.0.7 (12). Three marker genes (PF05958, PF06723 and PF07991) were excluded in the calculation because they either were absent or multi-copied in the closed genome HKCCA1288. The ANI between genomes was calculated using fastANI v1.3 (13) with the default parameters.

Protein-coding genes were predicted with the Prokka annotation pipeline v1.12 (14). Protein sequences were annotated using the online RAST (15) and KEGG server (16). They were further searched against the COG (17), Pfam (18), TIGRFAM (19) and CDD (20) databases, all of which were downloaded in February 2020. Proteins involved in amino acid biosynthesis were inferred using the online tool GapMind (21).

#### *1.3 Phylogenomic analysis*

To place the CHUG lineage into the phylogeny of the *Roseobacter* group, we performed a phylogenomic analysis based on 120 bacterial marker genes (22). Eight CHUG genomes, one newly sequenced reference genome (LSUCC0031), and 78 additional reference genomes from a previous study (23) were used for phylogenomic tree construction. Marker genes were each aligned at the amino acid sequence level using MAFFT v7.222 (24) and trimmed using trimAl v1.4.rev15 (25) with ‘-resoverlap 0.55 -seqoverlap 60’ options. The trimmed alignments were concatenated using a custom script (https://github.com/luolab-cuhk/CHUG-genome-reduction-project) to comprise a super-alignment with 456, 904 sites. The maximum likelihood (ML) phylogenomic tree was built using IQ-TREE v1.6.2 (26) with the ModelFinder (27) assigning the best substitution model, and a total of 1,000 ultrafast bootstrap replicates were sampled to assess the robustness of the phylogeny (28). The phylogeny was visualized using iTOL (29).

*1.4 Phylogenetic analysis based on 16S rRNA genes*

To demonstrate that the CHUG lineage represents a distinct phylogenetic branch in the *Roseobacter* group, we constructed an ML phylogeny based on 16S rRNA gene sequences of both cultured and uncultivated roseobacters following the same method mentioned above. The cultured roseobacters' 16S rRNA genes were extracted from the above-mentioned 87 genomes, and the uncultivated roseobacters' sequences were retrieved from a previous study (30). To broadly search uncultivated 16S rRNA gene sequences that are closely related to the cultured CHUG members, we built a preliminary 16S rRNA gene tree (data not shown) with all *Roseobacter* sequences from the SILVA database (31) and those from the CHUG genomes, and identified three SILVA sequences closely related to the CHUG isolates. These three sequences (JN119120, JQ197701 and GQ342302) were also included in the above-mentioned IQ-TREE phylogenetic analysis.

*1.5 Genome content analysis*

To identify the shared genomic content between CHUG and the previously reported pelagic *Roseobacter* cluster (PRC) (32), we reconstructed a dendrogram based on the presence and absence patterns of orthologous families. Orthologous gene families were identified using OrthoFinder v2.2.1 (33) with '-S diamond -M msa' options. The binary matrix of presence and absence pattern for each orthologous gene family was used for dendrogram construction using IQ-TREE v1.6.2 (26), and a total of 1,000 ultrafast bootstrap replicates were sampled to assess the robustness of the dendrogram (28). The dendrogram was visualized using iTOL (29).

To help understand the evolutionary process giving rise to the CHUG lineage, we reconstructed the genome content of the ancestral nodes related to CHUG, its sister lineage and their outgroups, and also inferred the gene gain and loss events using BadiRate v1.35 (34) with '-anc -bmodel FR -rmodel BDI -ep CSP' options. A pruned subtree of the phylogeny inferred based on 120 bacterial marker genes (Fig. 1A) and a table of gene count for each orthologous gene family predicted by OrthoFinder (33) were used as the inputs. The ancestral genome sizes were estimated based on the number of orthologous gene families under linear regression model ($R^2$ =0.97, $p < 0.01$) in R.

#### *1.6 Statistical analysis for genomic features*

The assembled genome size, gene number, coding density and GC content were obtained using CheckM v1.0.7 (12), then the estimated genome size was adjusted as (*assembled genome size*)/(*completeness* + *contamination*) (35). Pseudogenes were predicted following our recent study (36). The number of carbon atoms per amino-acid-residue side chain (C-ARSC), the number of nitrogen atoms per amino-acid-residue side chain (N-ARSC), amino acid usage, and codon usage were retrieved using custom scripts (https://github.com/luolab-cuhk/CHUG-genome-reduction-project). The number of orthologous families and the mean number of genes per orthologous family were calculated based on the gene families clustered by OrthoFinder v2.2.1 (33). Phylogenetic ANOVA analyses were performed to compare these genomic features between different *Roseobacter* lineages using the 'phylANOVA' function in the 'phytools' R package, which allows controlling for phylogenetic impact on these metrics (37).

Next, we identified genes that were either enriched or depleted in CHUG and other PRC members compared to other roseobacters. Briefly, the phylogenetic signal of functional genes (*coxL, pdo, sox*) or traits (light utilization) were checked with the 'phylosig' function of the 'phytools' R package (37). A strong phylogenetic signal was identified in the distribution of *pdo, sox* and the light utilization trait (lambda > 0.99, $p < 0.001$ for each), so the subsequent tests whether they were associated with a particular category (PRC or non-PRC) were controlled for evolutionary history using the 'binaryPGLMM' function in the 'ape' R package (38). On the other hand, there was no phylogenetic signal for the *coxL* gene (lambda < 0.01), so $\chi^2$ test was used to test whether it was associated with a particular category (PRC or non-PRC).

#### *1.7 Recruitment analysis using public metagenomic sequencing data*

To identify the global occurrence and activity of CHUG members, the *TARA* Ocean metagenomic and metatranscriptomic sequencing data (39–41) were downloaded and mapped to the genomes. Two additional metagenomic sequencing data sampled at the Red Sea (42) and the Kwangyang bay (43) were also collected because an increased abundance of CHUG members were identified in a preliminary analysis. These public sequencing data were quality trimmed with Trimmomatic v0.36 (8) with options 'SLIDINGWINDOW:4:15 MAXINFO:40:0.9 MINLEN:40' and were subsequently mapped to the 89 *Roseobacter* genomes using bowtie v2.3.2 (44) with the parameter '-very-sensitive-local' for a quick screening. Mapped reads were extracted using SAMtools v1.4.1 (45) and more precisely searched against the 89 genomes using BLASTN (46) with the setting '-evalue 1e-5 -perc_identity 95 -qcov_hsp_perc 80'. Thereby only those mapped reads that shared >95% identity and >80% coverage with a reference genome were kept for further calculations. The relative abundances of CHUG and other PRC members were represented using Reads Per

Kilobase per Million mapped reads (RPKM) and compared using the Wilcox test in the 'ggplot2' R package. The correlation analysis was performed using the 'rcorr' function in the 'Hmisc' R package (47), and the significance level was adjusted using stringent Bonferroni correction. These analyses were not performed for each CHUG genome individually because these genomes are closely related (95.4 ± 2.4% ANI) and some reads were equally mapped to multiple genomes.

*1.8 $d_R/d_C$ ratio calculation*

We calculated the ratio of radical nonsynonymous nucleotide substitutions per radical nonsynonymous site ($d_R$) versus conservative nonsynonymous nucleotide substitutions per conservative nonsynonymous site ($d_C$) following our previous protocol (48). Briefly, orthologous gene families from CHUG members and their sister group were compared to those from the outgroup, respectively. The 20 amino acids were categorized into three groups based on charge or six groups according to volume and polarity (48). For each method of categorization, a nonsynonymous nucleotide substitution was considered 'conservative' if it led to a within-group replacement of amino acids and 'radical' if it resulted in a between-group replacement of amino acids. The $d_R/d_C$ ratio was calculated using three methods: GC-corrections based on codon frequency, GC-corrections based on amino acid composition, and the traditional uncorrected method (49), all of which were implemented in RCCalculator (48).

*1.9 Vitamin $B_{12}$ assay*

To validate the vitamin $B_{12}$ auxotrophy in CHUG members, a growth assay was performed for the HKCCA1288 as the experimental CHUG strain and the model roseobacter strain *Ruegeria pomeroyi* DSS-3 (50) as the positive control. The starter culture was prepared in 2216 marine broth (Difco) prior to the growth experiments. In the vitamin $B_{12}$ assay, the defined marine ammonium mineral salts (MAMS) medium was supplemented with ribose (30 mM, Sigma) as the sole carbon source and with SL-10 trace metals solution (1 mL/L) (51). Vitamin mixture in the presence or absence of vitamin $B_{12}$ was added as described previously (52). Strains were cultivated in MAMS medium for 96 h, and samples were collected every 12 h for cell counting. The collected cells were stained with SYBR Green I (Life Technologies, USA) dye for 15 min, and cell numbers were counted using a flow cytometer (Guava EasyCyte Plus, MA, USA) equipped with a fluorescence detector. All experiments in this section were performed in triplicate.

### Text 2. Supplementary results

#### *2.1 Biased codon and amino acid usage*

As mentioned in the main paper, the GC content (Fig. 2C) was lower in both CHUG members and seven other PRC genomes compared to non-PRC members. We further investigated whether there was codon usage bias within these genomes for the 18 amino acids each encoded by more than one codon (Fig. S2). CHUG members tended to use more adenine/thymine (A/T) in the synonymous codons encoding 11 and 12 amino acids when compared to its sister group and the outgroup, respectively (Fig. S2). For example, among the four synonymous codons for proline (CCA, CCT, CCG, CCC), CCA and CCT were more frequently used in CHUG (20.9% and 23.7%, respectively) compared to its sister group (7.4% and 8.3%, respectively; $p < 0.05$ for each) and outgroup (7.1%, $p > 0.05$ for CCA; 7.5%, $p < 0.01$ for CCT). In contrast, the frequencies of the rest codons (CCG and CCC) were reduced in CHUG (22.6% and 32.6%, respectively) compared to its sister group (42.2% and 41.9%, respectively; $p < 0.05$ for each) and outgroup (45.8%, $p < 0.05$ for CCG; 39.4%, $p > 0.05$ for CCC).

Next, we investigated the amino acid usage frequency bias within the oligotrophic

CHUG and seven other PRC members (Fig. S3). The 20 amino acids can be divided into six groups based on their volume and polarity (Fig. S3) (53). Among the nonpolar and relatively small amino acids, CHUG members tended to use more isoleucine (5.7%) and less valine (6.8%) compared to the outgroup (4.9% for isoleucine and 7.3% for valine; $p < 0.05$). One possible explanation was that the codons for isoleucine (ATT, ATC and ATA) generally used less C and G and thus saved more nitrogen than the codons for valine (GTT, GTC, GTA, GTG) (54).

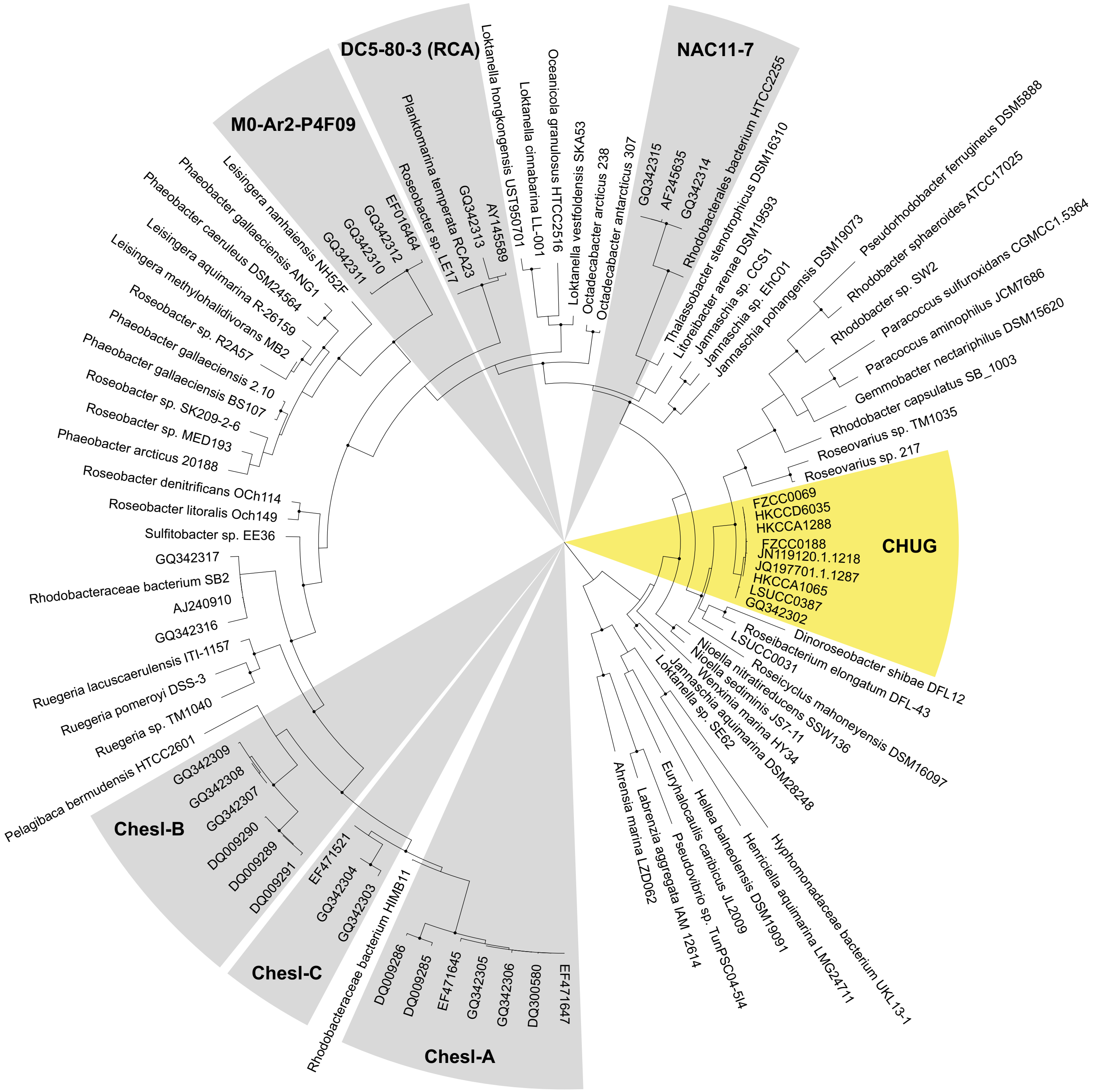


**Fig. S1.** The IQ-TREE maximum likelihood phylogenetic tree based on 16S rRNA gene sequences. The sequences include those extracted from the 87 genomes used in the present study, those collected from the SILVA database and clustered with CHUG, and those from the wild Roseobacter clusters named in a previous study (30). Solid circles in the phylogeny indicate nodes with bootstrap values >95%.

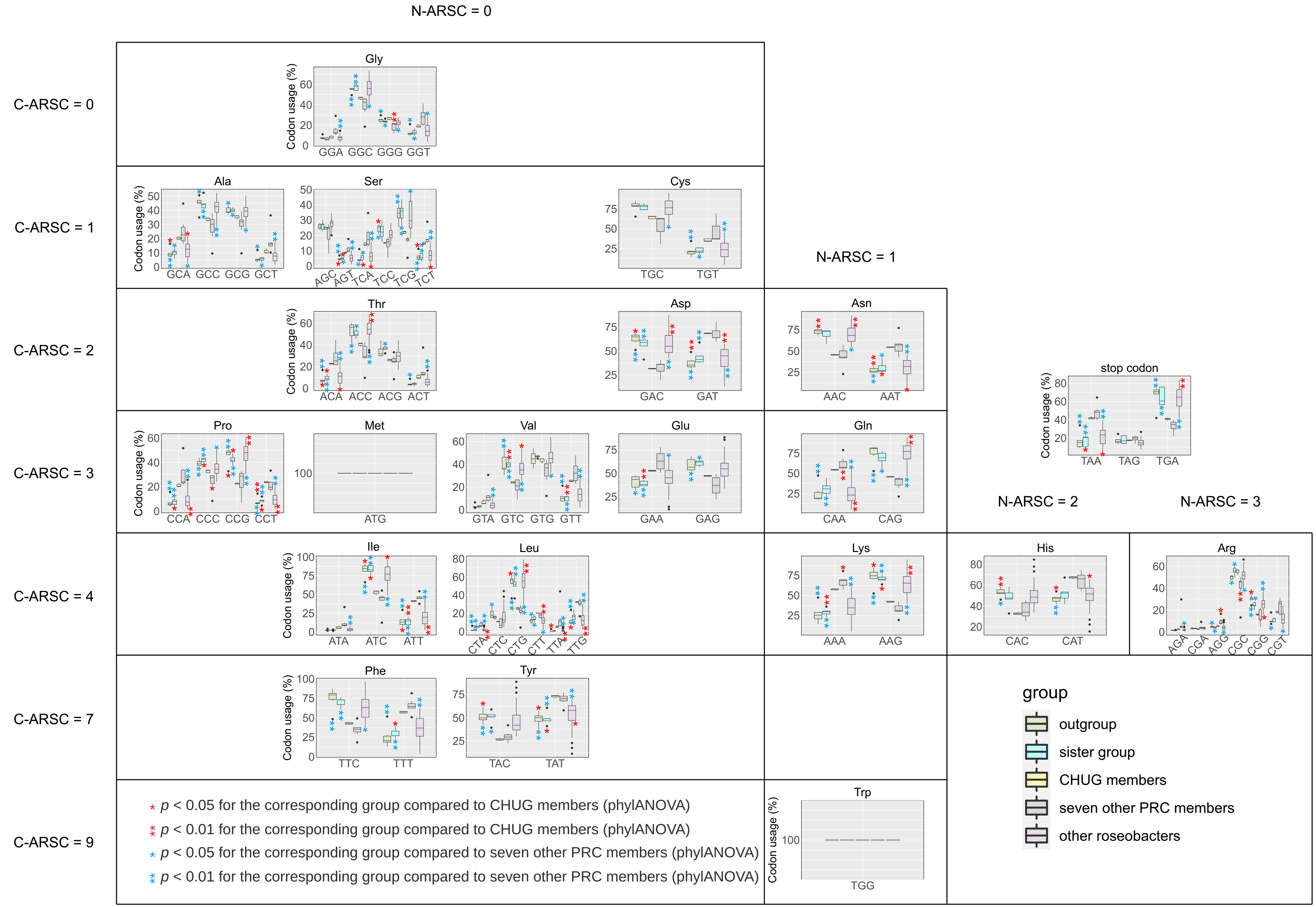


**Fig. S2.** Codon usage frequency between CHUG, its sister group, the outgroup, seven other PRC members, and other reference roseobacters. The significance level in the codon usage frequency between CHUG and other four groups are shown in red, while that between seven other PRC members and the remaining three groups are shown in blue. The markers * and ** denote $p < 0.05$ and $p < 0.01$ (phylANOVA analysis), respectively. The 20 amino acids are placed based on their number of carbon atoms per amino-acid-residue side chain (C-ARSC) and number of nitrogen atoms per amino-acid-residue side chain (N-ARSC).

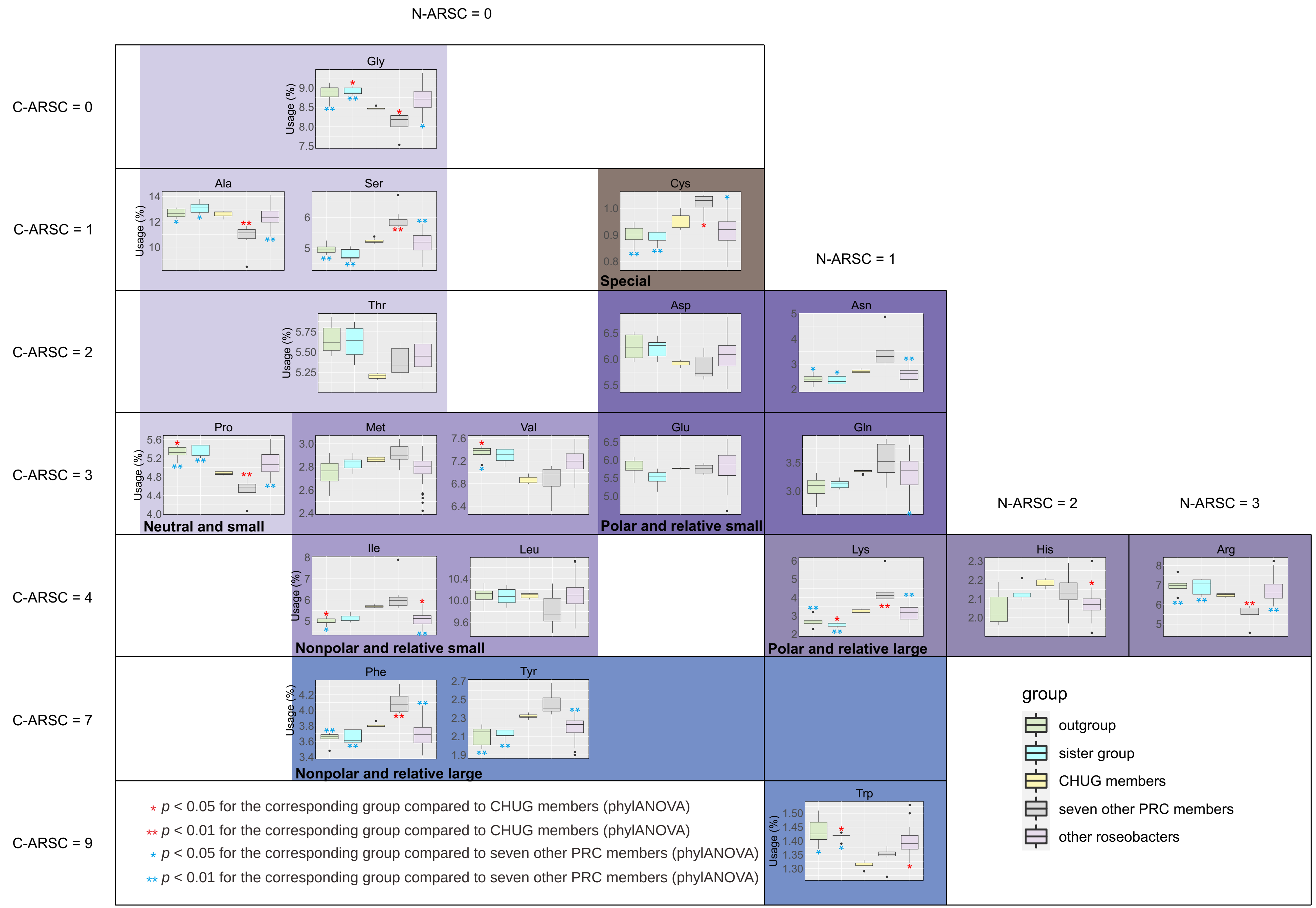


**Fig. S3.** Amino acid usage frequency between CHUG, its sister group, the outgroup, seven other PRC members, and other reference roseobacters. The significance level in the amino acid usage frequency between CHUG and other four groups are shown in red, while that between seven other PRC members and the remaining three groups are shown in blue. The markers * and ** denote $p < 0.05$ and $< 0.01$ (phylANOVA analysis), respectively. The amino acids are placed based on their number of carbon atoms per amino-acid-residue side chain (C-ARSC) and number of nitrogen atoms per amino-acid-residue side chain (N-ARSC). Six amino acid groups based on their volume and polarity were shaded with different color.

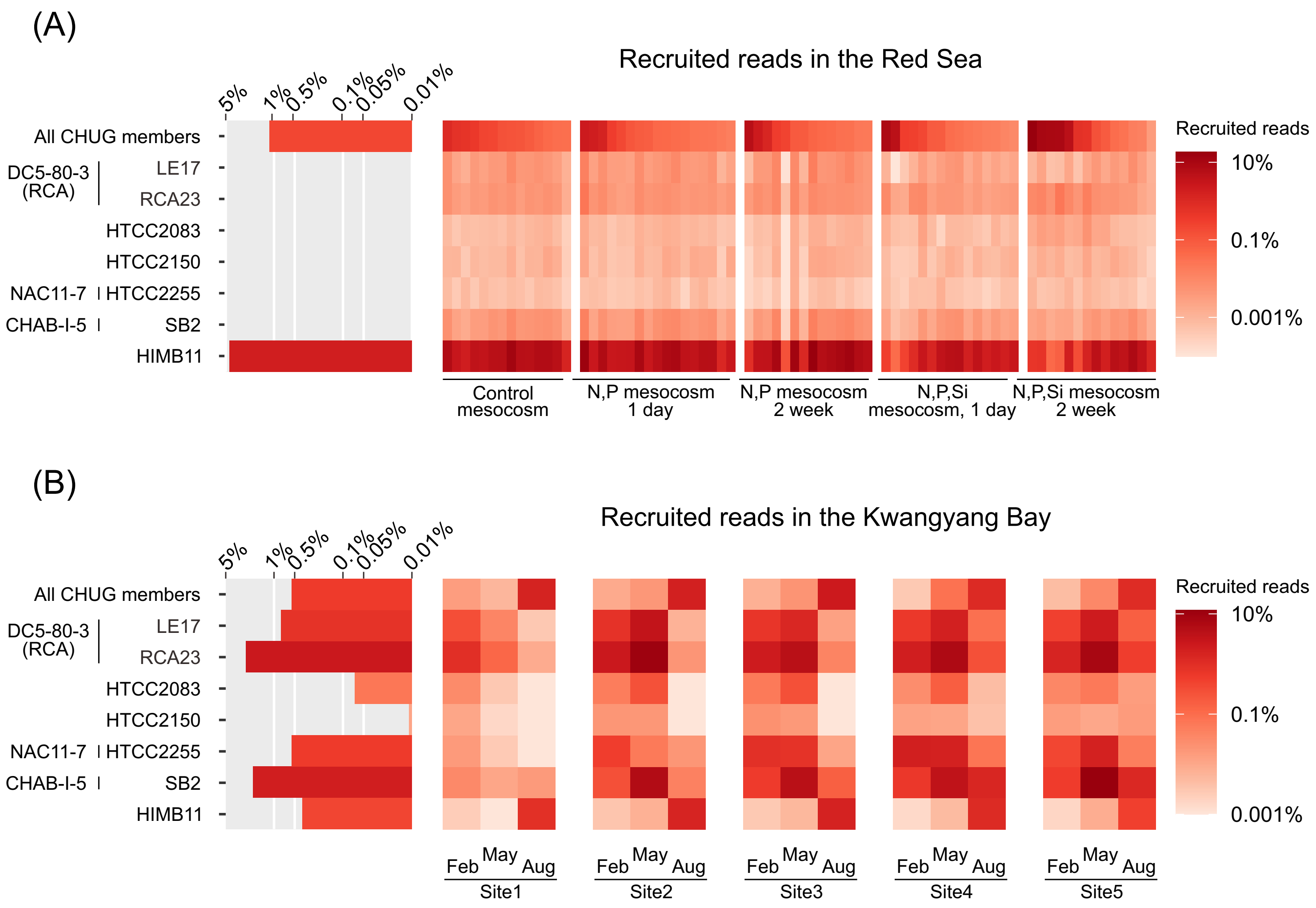


**Fig. S4.** The relative abundance of CHUG and other PRC members in the bacterioplankton communities based on recruitment analysis using the Red Sea (A) and Kwangyang bay (B) sequencing samples.

(A)

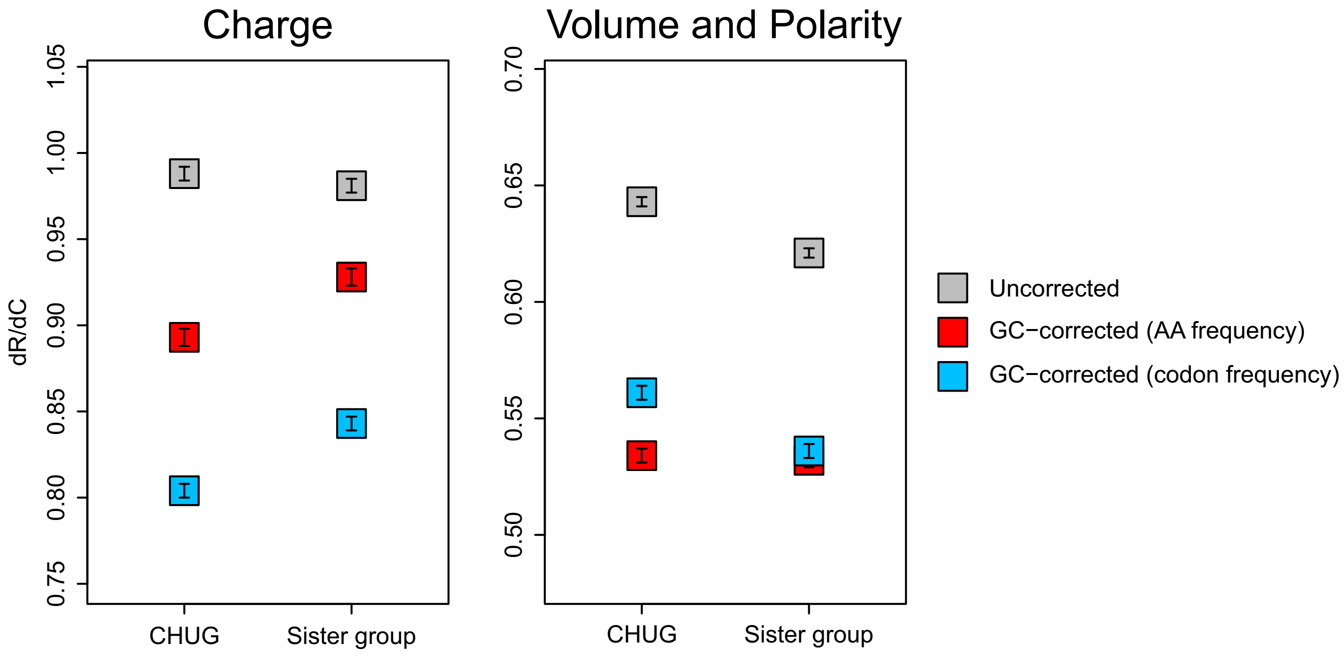


(B)

| Charge | Amino acid |
|---|---|
| Neutral | Alanine, Asparagine, Cysteine, Glutamine, Glycine, Isoleucine, Leucine, Methionine, Phenylalanine, Proline, Serine, Threonine, Tryptophan, Tyrosine, Valine |
| Positive | Arginine, Histidine, Lysine |
| Negative | Asparticacid Glutamicacid |

(C)

| Volume and polarity | Amino acid |
|---|---|
| special | Cysteine |
| neutral and small | Alanine, Glycine, Proline, Serine, Threonine |
| nonpolar and relative small | Isoleucine, Leucine, Methionine, Valine |
| nonpolar and relative large | Phenylalanine, Tryptophan, Tyrosine |
| polar and relative small | Asparticacid, Glutamicacid, Asparagine, Glutamine |
| polar and relative large | Histidine, Lysine, Arginine |

**Fig. S5.** (A) Analysis of the $d_R/d_C$ ratios of CHUG members and their sister group compared to their outgroup. These ratios are calculated based on the physicochemical classification of the 20 amino acids by charge (left) and by volume and polarity (right), respectively (48). Bars indicate one standard deviation of the mean. Results were obtained using RCCalculator (http://www.geneorder.com/RCCalculator) with GC-corrections based on codon frequency (blue square), amino acid (AA) composition (red square), and uncorrected method HON-NEW (grey square). (B) Classification of amino acids by charge. (C) Classification of amino acids by volume and polarity.

**References**


1. Mak SST, Gopalakrishnan S, Carøe C, Geng C, Liu S, Sinding M-HS et al. Comparative performance of the BGISEQ-500 vs Illumina HiSeq2500 sequencing platforms for palaeogenomic sequencing. Gigascience 2017; 6(8):1–13.

2. Lui W-Y, Yuen C-K, Li C, Wong WM, Lui P-Y, Lin C-H et al. SMRT sequencing revealed the diversity and characteristics of defective interfering RNAs in influenza A (H7N9) virus infection. Emerg Microbes Infect 2019; 8(1):662–74.

3. Connon SA, Giovannoni SJ. High-throughput methods for culturing microorganisms in very-low-nutrient media yield diverse new marine isolates. Appl Environ Microbiol 2002; 68(8):3878–85.

4. Song J, Oh H-M, Cho J-C. Improved culturability of SAR11 strains in dilution-to-extinction culturing from the East Sea, West Pacific Ocean. FEMS Microbiol Lett 2009:141–7.

5. Yang S-J, Kang I, Cho J-C. Expansion of cultured bacterial diversity by large-scale dilution-to-extinction culturing from a single seawater sample. Microb Ecol 2016:29–43.

6. Henson MW, Pitre DM, Weckhorst JL, Lanclos VC, Webber AT, Thrash JC. Artificial seawater media facilitate cultivating members of the microbial majority from the Gulf of Mexico. mSphere 2016; 1(2).

7. Henson MW, Lanclos VC, Pitre DM, Weckhorst JL, Lucchesi AM, Cheng C et al. Expanding the diversity of bacterioplankton isolates and modeling isolation efficacy with large-scale dilution-to-extinction cultivation. Appl Environ Microbiol 2020; 86(17).

8. Bolger AM, Lohse M, Usadel B. Trimmomatic: a flexible trimmer for Illumina sequence data. Bioinformatics 2014; 30(15):2114–20.

9. Bankevich A, Nurk S, Antipov D, Gurevich AA, Dvorkin M, Kulikov AS et al. SPAdes: a new genome assembly algorithm and its applications to single-cell sequencing. J Comput Biol 2012; 19(5):455–77.

10. Babraham Bioinformatics. FastQC: a quality control tool for high throughput sequence data. Cambridge, UK: Babraham Institute 2011.

11. Wick RR, Judd LM, Gorrie CL, Holt KE. Unicycler: Resolving bacterial genome

assemblies from short and long sequencing reads. PLoS Comput Biol 2017; 13(6):e1005595.

12. Parks DH, Imelfort M, Skennerton CT, Hugenholtz P, Tyson GW. CheckM: assessing the quality of microbial genomes recovered from isolates, single cells, and metagenomes. Genome Res 2015; 25(7):1043–55.

13. Jain C, Rodriguez-R LM, Phillippy AM, Konstantinidis KT, Aluru S. High throughput ANI analysis of 90K prokaryotic genomes reveals clear species boundaries. Nat Commun 2018; 9(1):5114.

14. Seemann T. Prokka: rapid prokaryotic genome annotation. Bioinformatics 2014; 30(14):2068–9.

15. Brettin T, Davis JJ, Disz T, Edwards RA, Gerdes S, Olsen GJ et al. RASTtk: a modular and extensible implementation of the RAST algorithm for building custom annotation pipelines and annotating batches of genomes. Sci Rep 2015; 5:8365.

16. Kanehisa M, Goto S. KEGG: kyoto encyclopedia of genes and genomes. Nucleic Acids Res 2000; 28(1):27–30.

17. Tatusov RL, Fedorova ND, Jackson JD, Jacobs AR, Kiryutin B, Koonin EV et al. The COG database: an updated version includes eukaryotes. BMC Bioinformatics 2003; 4:41.

18. Finn RD, Bateman A, Clements J, Coggill P, Eberhardt RY, Eddy SR et al. Pfam: the protein families database. Nucleic Acids Res 2014; 42(Database issue):D222-30.

19. Haft DH, Selengut JD, White O. The TIGRFAMs database of protein families. Nucleic Acids Res 2003; 31(1):371–3.

20. Marchler-Bauer A, Lu S, Anderson JB, Chitsaz F, Derbyshire MK, DeWeese-Scott C et al. CDD: a Conserved Domain Database for the functional annotation of proteins. Nucleic Acids Res 2011; 39(Database issue):D225-9.

21. Price M, Deutschbauer AM, Arkin AP. GapMind: Automated annotation of amino acid biosynthesis; 2019.

22. Parks DH, Chuvochina M, Waite DW, Rinke C, Skarshewski A, Chaumeil P-A et al. A standardized bacterial taxonomy based on genome phylogeny substantially revises the tree of life. Nat Biotechnol 2018; 36(10):996–1004.

23. Simon M, Scheuner C, Meier-Kolthoff JP, Brinkhoff T, Wagner-Döbler I, Ulbrich M et al. Phylogenomics of *Rhodobacteraceae* reveals evolutionary adaptation to marine and non-

marine habitats. ISME J 2017; 11(6):1483–99.

24. Katoh K, Standley DM. MAFFT multiple sequence alignment software version 7: improvements in performance and usability. Mol Biol Evol 2013; 30(4):772–80.

25. Capella-Gutiérrez S, Silla-Martínez JM, Gabaldón T. trimAl: a tool for automated alignment trimming in large-scale phylogenetic analyses. Bioinformatics 2009; 25(15):1972–3.

26. Nguyen L-T, Schmidt HA, Haeseler A von, Minh BQ. IQ-TREE: a fast and effective stochastic algorithm for estimating maximum-likelihood phylogenies. Mol Biol Evol 2015; 32(1):268–74.

27. Kalyaanamoorthy S, Minh BQ, Wong TKF, Haeseler A von, Jermiin LS. ModelFinder: fast model selection for accurate phylogenetic estimates. Nat Methods 2017; 14(6):587–9.

28. Hoang DT, Chernomor O, Haeseler A von, Minh BQ, Le Vinh S. UFBoot2: Improving the ultrafast bootstrap approximation. Mol Biol Evol 2018; 35(2):518–22.

29. Letunic I, Bork P. Interactive Tree Of Life (iTOL) v4: recent updates and new developments. Nucleic Acids Res 2019.

30. Buchan A, Hadden M, Suzuki MT. Development and application of quantitative-PCR tools for subgroups of the *Roseobacter* clade. Appl Environ Microbiol 2009; 75(23):7542–7.

31. Quast C, Pruesse E, Yilmaz P, Gerken J, Schweer T, Yarza P et al. The SILVA ribosomal RNA gene database project: improved data processing and web-based tools. Nucleic Acids Res 2013; 41(Database issue):D590-6.

32. Billerbeck S, Wemheuer B, Voget S, Poehlein A, Giebel H-A, Brinkhoff T et al. Biogeography and environmental genomics of the *Roseobacter*-affiliated pelagic CHAB-I-5 lineage. Nat Microbiol 2016; 1(7):16063.

33. Emms DM, Kelly S. OrthoFinder: phylogenetic orthology inference for comparative genomics. Genome Biol 2019; 20(1):238.

34. Librado P, Vieira FG, Rozas J. BadiRate: estimating family turnover rates by likelihood-based methods. Bioinformatics 2012; 28(2):279–81.

35. Parks DH, Rinke C, Chuvochina M, Chaumeil P-A, Woodcroft BJ, Evans PN et al. Recovery of nearly 8,000 metagenome-assembled genomes substantially expands the tree of life. Nat Microbiol 2017; 2(11):1533–42.

36. Chu X, Li S, Wang S, Luo D, Luo H. Gene loss through pseudogenization contributes to the ecological diversification of a generalist *Roseobacter* lineage. ISME J 2020.

37. Revell LJ. phytools: an R package for phylogenetic comparative biology (and other things). Methods in Ecology and Evolution 2012; 3(2):217–23.

38. Paradis E, Schliep K. ape 5.0: an environment for modern phylogenetics and evolutionary analyses in R. Bioinformatics 2019; 35(3):526–8.

39. Sunagawa S, Coelho LP, Chaffron S, Kultima JR, Labadie K, Salazar G et al. Ocean plankton. Structure and function of the global ocean microbiome. Science 2015; 348(6237):1261359.

40. Salazar G, Paoli L, Alberti A, Huerta-Cepas J, Ruscheweyh H-J, Cuenca M et al. Gene expression changes and community turnover differentially shape the global ocean metatranscriptome. Cell 2019; 179(5):1068-1083.e21.

41. Vargas C de, Audic S, Henry N, Decelle J, Mahé F, Logares R et al. Eukaryotic plankton diversity in the sunlit ocean. Science 2015; 348(6237):1261605.

42. Coello-Camba A, Diaz-Rua R, Duarte CM, Irigoien X, Pearman JK, Alam IS et al. Picocyanobacteria community and cyanophage infection responses to nutrient enrichment in a mesocosms experiment in oligotrophic waters. Front. Microbiol. 2020; 11.

43. Kim Y, Jeon J, Kwak MS, Kim GH, Koh I, Rho M. Photosynthetic functions of *Synechococcus* in the ocean microbiomes of diverse salinity and seasons. PLoS ONE 2018; 13(1):e0190266.

44. Langmead B, Salzberg SL. Fast gapped-read alignment with Bowtie 2. Nat Methods 2012; 9(4):357–9.

45. Li H, Handsaker B, Wysoker A, Fennell T, Ruan J, Homer N et al. The sequence alignment/map format and SAMtools. Bioinformatics 2009; 25(16):2078–9.

46. Altschul SF, Gish W, Miller W, Myers EW, Lipman DJ. Basic local alignment search tool. J Mol Biol 1990; 215(3):403–10.

47. Harrell Jr FE. Package ‘Hmisc’. CRAN2018 2019; 2019:235–6.

48. Luo H, Huang Y, Stepanauskas R, Tang J. Excess of non-conservative amino acid changes in marine bacterioplankton lineages with reduced genomes. Nat Microbiol 2017; 2:17091.

49. Zhang J. Rates of conservative and radical nonsynonymous nucleotide substitutions in mammalian nuclear genes. J Mol Evol 2000; 50(1):56–68.

50. Moran MA, Buchan A, González JM, Heidelberg JF, Whitman WB, Kiene RP et al. Genome sequence of *Silicibacter pomeroyi* reveals adaptations to the marine environment. Nature 2004; 432(7019):910–3.

51. Lidbury I, Kimberley G, Scanlan DJ, Murrell JC, Chen Y. Comparative genomics and mutagenesis analyses of choline metabolism in the marine *Roseobacter* clade. Environ Microbiol 2015; 17(12):5048–62.

52. Kanagawa T, Dazai M, Fukuoka S. Degradation of O,O-dimethyl phosphorodithioate by *Thiobacillus thioparus* TK-1 and Pseudomonas AK-2. Agricultural and Biological Chemistry 1982; 46(10):2571–8.

53. Miyata T, Miyazawa S, Yasunaga T. Two types of amino acid substitutions in protein evolution. J Mol Evol 1979; 12(3):219–36.

54. Bohlin J, Brynildsrud O, Vesth T, Skjerve E, Ussery DW. Amino acid usage is asymmetrically biased in AT- and GC-rich microbial genomes. PLoS ONE 2013; 8(7):e69878.
